# Stability of N-type inactivation and the coupling between N-type and C-type inactivation in the *Aplysia* Kv1 channel

**DOI:** 10.1101/2024.02.19.581087

**Authors:** Tokunari Iwamuro, Kazuki Itohara, Yasuo Furukawa

## Abstract

The voltage-dependent potassium channels (Kv channels) show several different types of inactivation. N-type inactivation is a fast inactivating mechanism, which is essentially an open pore blockade by the amino-terminal structure of the channel itself or the auxiliary subunit. There are several functionally discriminatable slow inactivation (C-type, P-type, U-type), the mechanism of which is supposed to include rearrangement of the pore region. In some Kv1 channels, the actual inactivation is brought about by coupling of N-type and C-type inactivation (N-C coupling). In the present study, we focused on the N-C coupling of the *Aplysia* Kv1 channel (AKv1). AKv1 shows a robust N-type inactivation, but its recovery is almost thoroughly from C-type inactivated state owing to the efficient N-C coupling. In the I8Q mutant of AKv1, we found that the inactivation as well as its recovery showed two kinetic components apparently correspond to N-type and C-type inactivation. Also, the cumulative inactivation which depends on N-type mechanism in AKv1 was hindered in I8Q, suggesting that N-type inactivation of I8Q is less stable. We also found that Zn^2+^ specifically accelerates C-type inactivation of AKv1, and that H382 in the pore turret is involved in the Zn^2+^ binding. Because the region around Ile^8^ (I8) in AKv1 has been suggested to be involved in the pre-block binding of the amino-terminal structure, our results strengthen a hypothesis that the stability of the pre-block state is important for stable N-type inactivation as well as the N-C coupling in the Kv1 channel inactivation.

## Introduction

Electrical signaling is a universal mechanism for communication in the biological system. The electrical signaling is important for the intercellular communication as well as the intracellular signaling, and depends on the ion transport across the membrane by specialized families of membrane proteins called ion channels [21, 45]. Since monumental series of studies by Hodgkin and Huxley in 1952 [22, 23, 24, 25], the mechanisms for the activation and inactivation of voltage-gated ion channels continue to attract physiologists and biophysicists [21]. Among the numerous ion channels, voltage-gated potassium channels (Kv channels) are vital for repolarization of action potentials and determine electrical excitability of the cells.

From the late 1980s, the molecular mechanisms of inactivation in Kv channels have been resolved by using *Drosophila Shaker* channel (*Drosophila* Kv1 channel) as a model system. A fast inactivation of *Drosophila Shaker* channel has been extensively examined and the inactivation (N-type inactivation) is found to be an open channel block by the amino-terminal region of the channel itself [10, 27, 71]. The mechanism is well explained by the ball and chain model originally proposed for the inactivation of Na^+^ channel [3]. N-type inactivation of *Drosophila Shaker* channel is absent in the amino-terminal deletion mutant of the channel (Sh-IR) and the apparent inactivation can be reproduced by a peptide block in Sh-IR [27, 46, 47, 71]. In later works done in mammalian Kv1 channels, the auxiliary subunit for Kv1 channels (Kv*β*) is shown to act as a pore blocking particle in intrinsically non-inactivating (or slowly inactivating) Kv1 channels [58]. Although the N-terminal docked structure of Kv1 channel is not yet available (but see, [13]), detailed structural inferences of N-type inactivation in Kv1 channels have been documented [20, 54, 66, 67, 73]. Although N-type inactivation can be approximated by a single-step pore block, it is actually multistep process [54, 73]. Earlier step for the pore block (the preblock) seems to be a binding of an intermediate region of the amino-terminal structure to the T1 domain beneath the transmembrane domain of Kv1 channels [20, 53, 73].

Sh-IR still shows much slower inactivation named C-type inactivation [28]. C-type inactivation is more prevalent mechanism for the inactivation in many Kv channels which lack N-type mechanism and the retardation of the inactivation by external K^+^ and TEA is a hallmark of C-type inactivation [7, 19, 35, 40]. Molecular mechanisms of the C-type inactivation of Kv channels are also extensively studied [26, 32, 37, 39, 48, 69]. Although recent structural analyses show that a dilation of the selectivity filter of the pore is a key step of C-type inactivation in Kv channels [49, 57, 64, 65], the mechanism is challenged by analysis of C-type inactivation in the constitutively active mutants of *Drosophila Shaker* channel [8]. Although C-type inactivation is the most documented mechanism for the slow inactivation of Kv channels, the slow inactivation is rather complex and other functionally separable mechanisms (P-type inactivation, U-type inactivation) are also known in some Kv channels [9, 33].

In some Kv1 channels which possess both N-type and C-type mechanisms, the inactivation of the channels is not simply a sum of two mechanisms but actually a coupled process, in which the C-type inactivation coupled to N-type inactivation is much faster [28] (hereafter, we call this coupling N-C coupling). Baukrowitz and Yellen have shown clearly that the N-C coupling is essential for the frequency dependent cumulative inactivation in *Drosophila Shaker* channel, and have estimated that *∼*50% of the recovery from inactivation at the holding potential of -80 mV following a 20 msec depolarizing pulse is from the C-type inactivated state [4]. Similar but more efficient N-C coupling is seen in mammalian Kv1.4 and *Aplysia* Kv1 channel (AKv1), which are supposed to recover almost exclusively from the C-type inactivated state [54, 56]. In considering the N-C coupling, a point mutant of AKv1 (I8Q) is quite interesting because I8Q shows a large stationary current in high K^+^ condition in spite of the fact that the kinetics of N-type inactivation is similar to the wild-type AKv1 [51].

In the present study, we compared the inactivation of AKv1 and I8Q in different K^+^ condition and tested the effects of some agents which are known to affect C-type inactivation in some Kv channels. We found that I8Q can enter the N-type inactivated state as in AKv1 but the following entry into C-type inactivated state is limited because of the low efficient N-C coupling. The present results suggest that a key to the efficient N-C coupling is the stability of the pre-block state in N-type inactivation.

## Materials and Methods

### Ethics

All animal experiments were approved by the Hiroshima University Animal Research Committee (No. G20-1), and performed in accordance with the guidelines for the Japanese Association of Laboratory Animal Science and the Animal Experimentation of Hiroshima University.

### Channels

An *Aplysia* Kv1 channel (AKv1) [52] was used in the present study. The plasmids containing the wild-type AKv1 (pSPAK01) and the amino-terminal deletion mutant of AKv1 (ΔN, pSPAK01ΔN, the 2nd–61th amino acids were removed) as well as the plasmids containing mutants (pSPAK01-A378E, pSPAK01-D379P, pSPAK01ΔN-A378E) were described previously [15, 60]. I8Q mutant of AKv1 (pSPAK01-I8Q) as well as H382Q mutants (pSPAK01-H382Q, pSPAK01ΔN-H382Q) were made by Quick Change protocol (Agilent Technologies, Santa Clara, CA, USA). The double mutant channels (I8Q-A378E, I8Q-H382Q) were made by swapping StuI–BamHI fragment of pSPAK01-I8Q with the corresponding region of pSPAK01-A378E or pSPAK01ΔN-H382Q. In the following, the wild-type AKv1 is simply denoted as AKv1.

### Oocyte expression for electrophysiological recording

AKv1 as well as the mutant channels were expressed in *Xenopus laevis* oocytes and the oocytes were cultured as described previously [52]. Briefly, a part of ovary was dissected out from a frog which was anesthetized in 0.15% MS-222 (Sigma-Aldrich, St. Louis, MO, USA). The frog was temporarily maintained in a recovery tank after suturing the incision. After confirming the postoperative recovery, the frog was returned to a home tank. The ovary was digested by 2% collagenase (Wako Chemicals, Osaka, Japan) dissolved in OR2 medium (in mM: NaCl 82.5, KCl 2, MgCl_2_ 1, HEPES 10, pH 7.5) for 1–2 hours. Dissociated oocytes were collected and washed by ND96 (in mM: NaCl 96, KCl 2, CaCl_2_ 1.8, MgCl_2_ 1, HEPES 10, pH 7.5). Stage V–VI oocytes were selected and incubated at 18 *^◦^*C in ND96 for 5 to 24 hours. Plasmids were digested with EcoRI and cRNA was synthesized by SP6 RNA polymerase using mMESSAGE mMACHINE SP6 (Thermo Fisher Scientific, Waltham, MA, USA). Immediately before the injection into oocytes, cRNA was diluted appropriately with RNase-free water to control the expression level. 50nl of cRNA solution was routinely injected into the animal hemisphere of oocyte and the cRNA-injected oocytes were further incubated for 2–5 days before electrophysiological recording.

### Electrophysiological recording

Membrane currents of oocytes were measured by two-electrode voltage clamp by using OC-725C (Warner Instruments LLC., Hamden, CT, USA) as described previously [60]. An oocyte was placed in a recording chamber (*∼*100 *µ*l), and continuously perfused with ND96 by gravity (*∼*2 ml/min). The microelectrodes of 0.5–3 MΩ filled with 1M KCl were used for both potential measurement and current injection. Holding potential was –80 mV throughout in the present study. Membrane potential was controlled by CLAMPEX in the pCLAMP package (ver. 6, Axon Instruments, USA) and the digitized currents were stored in the hard disk for later analysis. Standard external solution was ND96. We also used a high K^+^ solution and a solution containing tetraethylammonium chloride instead of NaCl (TEA96). The composition of the solutions was as follows (mM): high K^+^ solution, KCl 98, CaCl_2_ 1.8, MgCl_2_ 1, HEPES 10 (pH 7.5); TEA96, TEA-Cl 96, KCl 2, CaCl_2_ 1.8, MgCl_2_ 1, HEPES 10 (pH 7.5). The solution containing appropriate amount of TEA was made by mixing ND96 and TEA96. We also examined the effect of Zn^2+^ on the inactivation of the channels. ZnCl_2_ was directly added to ND96 or high K^+^ solution.

In some experiments, the gating currents of AKv1 and some mutants were recorded by using non-conducting mutants (AKv1-W391F, I8Q-W391F, ΔN-W391F) made by Quick Change protocol (Agilent Technologies, Santa Clara, CA, USA). These non-conducting mutants correspond to a well-studied W434F mutant of *Drosophila Shaker* channel [50, 59, 69]. The gating currents were measured by cut-open vaseline gap voltage clamp (COVC) by using CA-1B (Dagan Corporation, Minneapolis, MN, USA) as described previously [62, 63]. Briefly, the top and middle compartments of the COVC chamber were filled with ND96 and the bottom compartment was filled with K-MES (in mM: KOH 100, EGTA 10, HEPES 10, pH was titrated by methansulfonic acid to 7.4). The oocyte membrane in the bottom compartment was permeabilized by 0.3–0.4% saponin or cut by scissors to establish electrical access to the inside of oocyte. The oocyte was internally dialyzed with K-MES via a glass pipette (*∼*200 *µ*m in diameter) connected to a syringe pump (flow rate was 2 *µ*l/min). The membrane potential of the oocyte was measured by a microelectrode of 0.2–1 MΩ filled with 0.5 M NaCl and controlled by CLAMPEX in the pCLAMP package (ver. 10, Molecular Device, San Jose, CA, USA). Holding potential was –80 mV and the leak currents were removed by P/4 or P/6 protocol (the holding potential for the subtraction pulses was -140 mV). The top compartment which isolates a domed membrane of the animal hemisphere of oocyte was continuously perfused with ND96 by gravity (*∼*1 ml/min).

All experiments were carried out at room temperature (20–25 *^◦^*C).

### Data analysis

Digitized data were analyzed with CLAMPFIT (Ver. 6, Axon Instruments; Ver. 10, Molecular Device) and Origin (Ver. 6, 8 or 10, Originlab, Northampton, MA, USA).

The steady state activation and inactivation of the channels were examined by conventional method as described previously [30]. The activation and inactivation curves were fitted with a Boltzmann equation as follows,

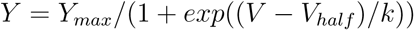

Y*_max_*, V*_half_*, and k are an estimated maximum value, half activation (or inactivation) voltage, and a slope factor, respectively.

The current decay in response to a depolarizing pulse was fitted with single or double exponential function as follows,

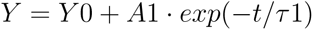

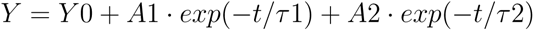

A1 or A2 is the amplitude of the inactivating component, *τ* 1 or *τ* 2 is the time constant of inactivation and Y0 is a remaining constant component.

The recovery from the inactivation as well as the cumulative inactivation was examined by a conventional two pulse protocol. The prepulse (1P, +40mV with variable duration) was followed by the test pulse (2P, +40mV, 20–40msec) with a variable inter-pulse interval and the peak currents (I*_peak_*) during 1P and 2P were measured. The inter-pulse potential was the same to the holding potential unless otherwise described. In each experiment, the current ratio (I*_peak_* in 2P / I*_peak_* in 1P) was plotted against the inter-pulse interval and the time course of recovery was fitted with exponential function as described above. In this case, Y0 means a size of fully recovered current, A1 or A2 means the size of recovering component from inactivation (a sign of A1 or A2 becomes minus in the recovery experiments), and *τ* 1 or *τ* 2 means the time constant for the recovery time course. After the fitting, the current ratio was normalized by Y0 in each experiment to make the recovery time course of pooled data shown in figures. The initial level of the normalized recovery (”init” in Tables) which indicates the proportion of activatable channels after the prepulse was estimated by 1 *− |A*1*|* for single exponential fitting or 1 *−* (*|A*1*|* + *|A*2*|*) for double exponential fitting.

The ON gating charge (Q*_on_*) was obtained by integrating the ON gating current during the pulse. The Q*_on_*-voltage relationship was analyzed by fitting to the Boltzmann equation as described above. The recovery time course of Q*_on_* was approximated by a single exponential function as described above. After each fitting, Y0 and A1 were normalized by Q*_on_* at 1st pulse to make the recovery time course of pooled data.

The results are presented as mean*±*SD. The statistical significance between two groups was estimated by Welch’s t-test. Multiple comparison among more than two groups was carried out by Dunnett’s test. Both tests were done by using R [55]. *p*-value of 0.05 was used as a critical value to reject the null-hypothesis.

## Results

### N-type inactivation of AKv1 and I8Q in ND96 and in high K^+^ solution

The functional property of N-type inactivation of AKv1 as well as its molecular mechanism are rather well characterized [14, 16, 51, 52, 53, 54, 60]. In the present study, we first reexamined the steady state activation and inactivation of AKv1, I8Q and ΔN and estimated the half activation/inactivation voltage (V*_half_*) and the slope factor (k). V*_half_* and k for the activation in ND96 were as follows (mV): AKv1, 0.9*±*8.3 & -6.3*±*3.8 (n=7); I8Q, 4.6*±*3.0 & -5.4*±*2.3 (n=9); ΔN, 6.7*±*1.6 & -6.0*±*0.8 (n=4). V*_half_* and k for the inactivation in ND96 were as follows (mV): AKv1, -16.0*±*5.7 & 2.3*±*0.3 (n=12); I8Q, -10.7*±*3.7 & 3.6*±*1.5 (n=13); ΔN, -7.4*±*0.7 & 3.2*±*0.5 (n=4). The parameters were in similar range to those described previously [15, 51, 52]. V*_half_* for either activation or inactivation of I8Q as well as ΔN was slightly shifted to more depolarized range compared to AKv1 as documented previously [51]. Fig. 1a shows families of the AKv1 currents in ND96 as well as in high K^+^ solution which is known to depress C-type inactivation [35, 40]. A series of depolarizing pulses of 1 sec were applied from the holding potential of -80mV every 20 sec and the current traces from -20 to +60 mV were superimposed. In response to the depolarizing step exceeding -20mV, the transient K^+^ current with little steady component was observed. The current decay was well approximated by a single exponential in ND96 as well as in high K^+^ solution (see insets of Fig. 1a). The time constant of inactivation (*τ_inacti_*) was voltage-independent at the membrane potential of more than +20 mV and *∼*20–30 msec (Fig. 1b). The appearance of inactivation was rarely affected by high K^+^ solution and *τ_inacti_*at +40 mV was similar in either ND96 or high K^+^ (see Table 1). Small steady current during the 1 sec pulse seemed to be slightly larger in high K^+^ solution (see Fig. 1a, Table 1), implying that the inactivated state of AKv1 is slightly less stable in high K^+^ condition.

**Fig. 1.**
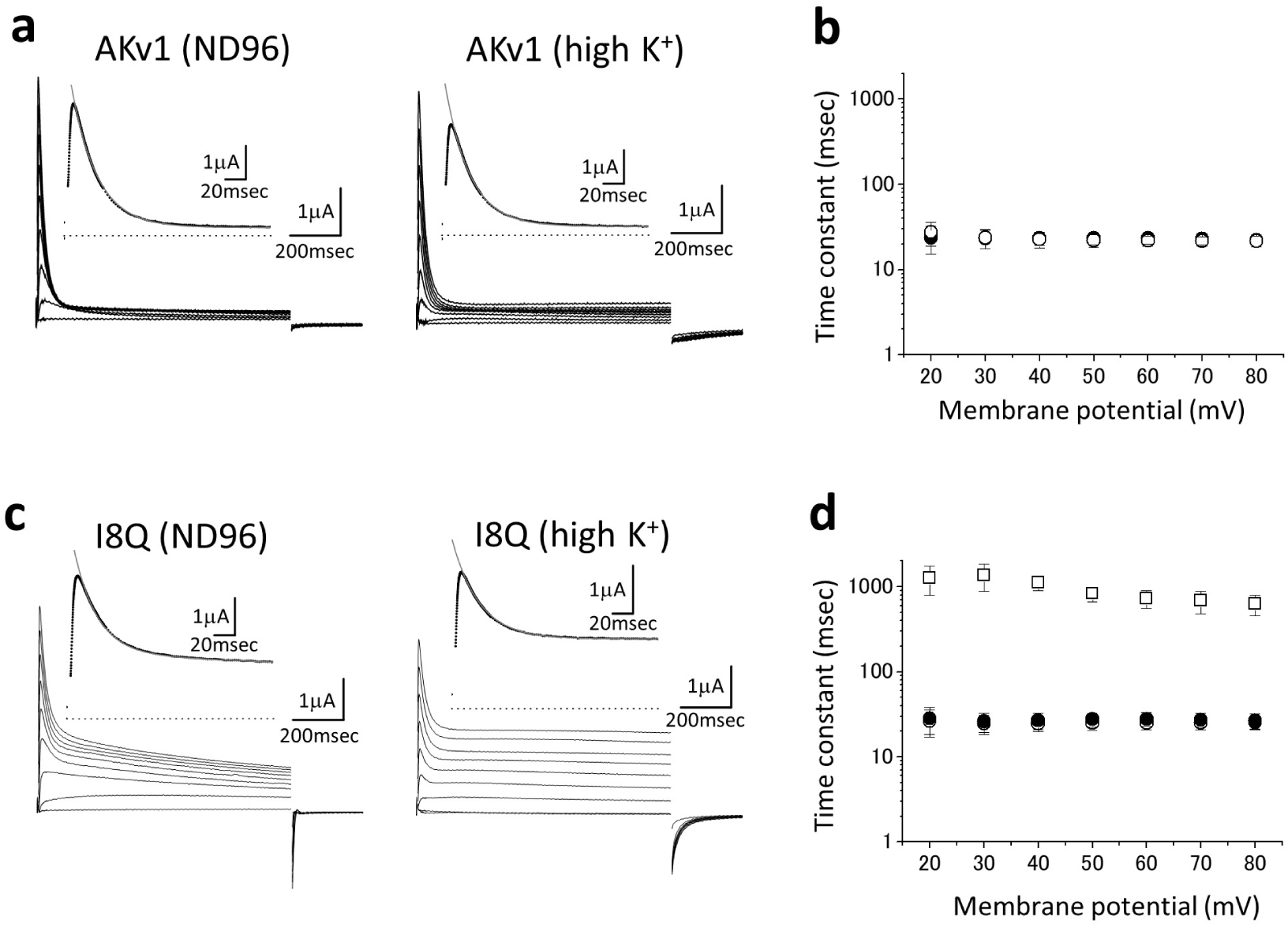
Comparison of the inactivation of AKv1 and I8Q. **a:** The inactivation of AKv1. Families of AKv1 currents in ND96 and in high K^+^ solution are shown. Step depolarizations of 1 sec from -20 to +60 mV were applied. The holding potential was -80mV. Insets show the time-expanded current traces at +40 mV and dotted lines indicate the zero current level. The current decay of AKv1 was approximated by a single exponential (thin line). The time constant (*τ_inacti_*) was 24.7 msec in ND96 and 23.8 msec in high K^+^. **b:** The relationship between *τ_inacti_*and the membrane potential in AKv1. In this and all other figures, symbols and error bars indicate mean*±*SD. Open circle (ND96), Filled circle (high K^+^). Note different symbols overlap mostly. **c:** The inactivation of I8Q. Families of I8Q currents in ND96 and in high K^+^ solution are shown. Insets show the time-expanded current traces at +40 mV as in **a**. Two time constants (fast and slow *τ_inacti_*) were required to fit the current decay of I8Q in ND96. The fast *τ_inacti_* and slow *τ_inacti_* in ND96 were 22.4 msec and 883.6 msec, respectively. The current decay of I8Q in high K^+^ solution was approximated by a single exponential and *τ_inacti_*was 26.2 msec. **d:** The relationship between *τ_inacti_*and the membrane potential in I8Q. Open circle (fast *τ_inacti_* in ND96), Square (slow *τ_inacti_*in ND96), Filled circle (*τ_inacti_*in high K^+^). Note *τ_inacti_*in high K^+^ solution is indistinguishable to the fast *τ_inacti_*in ND96 and *τ_inacti_*of AKv1.

**Table 1.** Inactivation of AKv1 and the I8Q mutants.

| channel | solution | $\tau 1$ | $p$ | $\tau 2$ | $p$ | A2/(A1+A2) | $p$ | st/peak | $p$ | n |
| --- | --- | --- | --- | --- | --- | --- | --- | --- | --- | --- |
| AKv1 | ND96 | 21.9±4.3 | – | NA | – | NA | NA | 0.078±0.045 | – | 18 |
| | high K<sup>+</sup> | 23.2±4.1 | 0.804 | NA | – | NA | NA | 0.135±0.076 | 0.015 | 13 |
| | 77mM TEA | 24.5±2.9 | 0.629 | NA | – | NA | NA | 0.01±0.01 | 0.087 | 4 |
| | 100μM Zn<sup>2+</sup> | 22.6±2.7 | 0.994 | NA | – | NA | NA | 0.031±0.011 | 0.245 | 5 |
| | 300μM Zn<sup>2+</sup> | 21.6±2.5 | 0.999 | NA | – | NA | NA | 0.039±0.014 | 0.420 | 5 |
| I8Q | ND96 | 24.2±4.2 | – | 964.2±119.9 | – | 0.398±0.046 | – | 0.232±0.077 | – | 28 |
| | high K<sup>+</sup> | 27.8±5.1 | 0.099 | NA | – | NA | NA | 0.512±0.071 | <1e-04 | 12 |
| | 77mM TEA | 27.3±5.1 | 0.577 | 987.8±258.7 | 0.972 | 0.447±0.148 | 0.331 | 0.26±0.08 | 0.993 | 4 |
| | 100μM Zn<sup>2+</sup> | 28.0±3.1 | 0.092 | 603.8±69.0 | <1e-05 | 0.47±0.052 | 0.004 | 0.31±0.258 | 0.267 | 11 |
| | 300μM Zn<sup>2+</sup> | 28.6±6.2 | 0.042 | 315.8±48.5 | <1e-05 | 0.512±0.053 | <1e-04 | 0.054±0.009 | 0.001 | 10 |
| I8Q-A378E | ND96 | 27.1±4.0 | – | 1465.3±689.4 | – | 0.214±0.082 | – | 0.551±0.07 | – | 9 |
| | high K<sup>+</sup> | 25.5±1.2 | 0.770 | 3354.9±1178.5 | 0.001 | 0.561±0.112 | <1e-04 | 0.665±0.046 | 0.012 | 6 |
| | 300μM Zn<sup>2+</sup> | 35.1±7.8 | 0.013 | 1941.1±376.0 | 0.501 | 0.542±0.128 | <1e-04 | 0.579±0.089 | 0.708 | 5 |
| I8Q-H382Q | ND96 | 25.5±3.6 | – | 356.5±54.1 | – | 0.306±0.038 | – | 0.331±0.054 | – | 11 |
| | high K<sup>+</sup> | 23.6±1.2 | 0.364 | 1389.9±1105.9 | 0.001 | 0.222±0.114 | 0.035 | 0.561±0.063 | <1e-04 | 5 |
| | 300μM Zn<sup>2+</sup> | 27.0±1.6 | 0.390 | 324.5±16.3 | 0.986 | 0.328±0.038 | 0.669 | 0.344±0.021 | 0.775 | 8 |

The inactivation (the current decay) was approximated by single or double exponential function.
$\tau 1$ , $\tau 2$ : fast ( $\tau 1$ ) and slow ( $\tau 2$ ) time constants (msec)
A2/(A1+A2): the relative amplitude of the slow inactivating component
st/peak: the relative amplitude of the current at the end of 1 sec pulse normalized by the peak amplitude
n: the number of tested oocytes
$p$ : $p$ -values obtained by Dunnett's test between ND96 and other solutions

I8Q is a point mutant of the amino-terminal subregion of AKv1 which is called the inactivation proximal (IP) region [51]. The macroscopic inactivation rate of I8Q is described to be similar to that of AKv1 in high K^+^ solution but the fractional amplitude of the inactivating component is much smaller [51]. Because the inactivation of I8Q in low K^+^ condition had not been documented, we examined the inactivation of I8Q in ND96 (2 mM K^+^) first and compared the results to those obtained in high K^+^ solution. In ND96, the voltage-dependent activation of I8Q was similar to AKv1 except for a slight depolarizing shift of the voltage-dependency as described above. The inactivation of I8Q showed two components in ND96 (Fig. 1c). The time constant of the fast component was *∼*20–30 msec and that of the slow component was *∼*1 sec (Fig. 1d, Table 1). As described previously [51], the inactivation of I8Q in high K^+^ solution was well approximated by a single exponential with a large steady component (Fig. 1c and 1d). The *τ_inacti_* of I8Q in high K^+^ solution was similar to the fast *τ_inacti_*in ND96 (Table 1).

### Comparison of the recovery from inactivation between AKv1 and I8Q

Compared to the fast inactivation, the recovery from inactivation of AKv1 is quite slow in ND96 [14, 52]. The time course of recovery from inactivation in AKv1 is similar to the recovery from C-type inactivation observed in the amino-terminal deletion mutant, suggesting that N-type inactivation of AKv1 is quite efficiently coupled to C-type inactivation and that the rate-limiting step of the recovery is a recovery from C-type inactivated state [54].

We compared the recovery from inactivation between AKv1 and I8Q in ND96 by a conventional two pulse protocol. We used a prepulse (1P, +40 mV) of variable duration (50–1000 msec). Following the prepulse, the membrane potential was returned to the holding potential of -80 mV for some time, and a test pulse (2P, +40 mV and 40 msec duration) was applied to assess the recovery from inactivation. In Fig. 2a, the AKv1 currents in response to the test pulses after the prepulse of 50 or 1000 msec with variable inter-pulse intervals at -80 mV are shown. The recovery from inactivation was analyzed by plotting the peak current at the test pulse against the inter-pulse interval. Fig. 2b compares the recovery time course of AKv1 after the prepulse of 50–1000 msec duration. In the present experiments, the shortest inter-pulse interval was 10 msec and the AKv1 current measured 10 msec after a 1 sec prepulse was less than a few percent of the peak current during the prepulse (see ”init” in Table 2). The current measured 10 msec after the prepulse was slightly larger when short prepulse was used (Table 2). The recovery time course after a long prepulse (500 or 1000 msec) was well fitted by a single exponential function with a time constant of *∼*3–4 sec, suggesting that a single slow rate limiting step dominates the recovery process (Table 2). Although a faster recovery process with the time constant of several hundred msec was noticeable in addition to the slow one when shorter prepulse was used (see Table 2), the slow component was still dominant (*>*70%). Fig. 2c shows the examples of recovery data in I8Q obtained by the similar pulse protocol. The recovery time course in I8Q was approximated by double exponential function (Fig. 2d, Table 2). The values of fast and slow *τ_rec_*(50–60 msec and 2-3 sec) were little affected by the duration of prepulse, but the relative amplitude of the slow component was clearly increased by lengthening the prepulse duration (Table 2). The results are consistent with a notion that I8Q recovers at least from two different inactivated states, and that the longer prepulse push more channels into the inactivated state from which the recovery is slow.

**Fig. 2.**
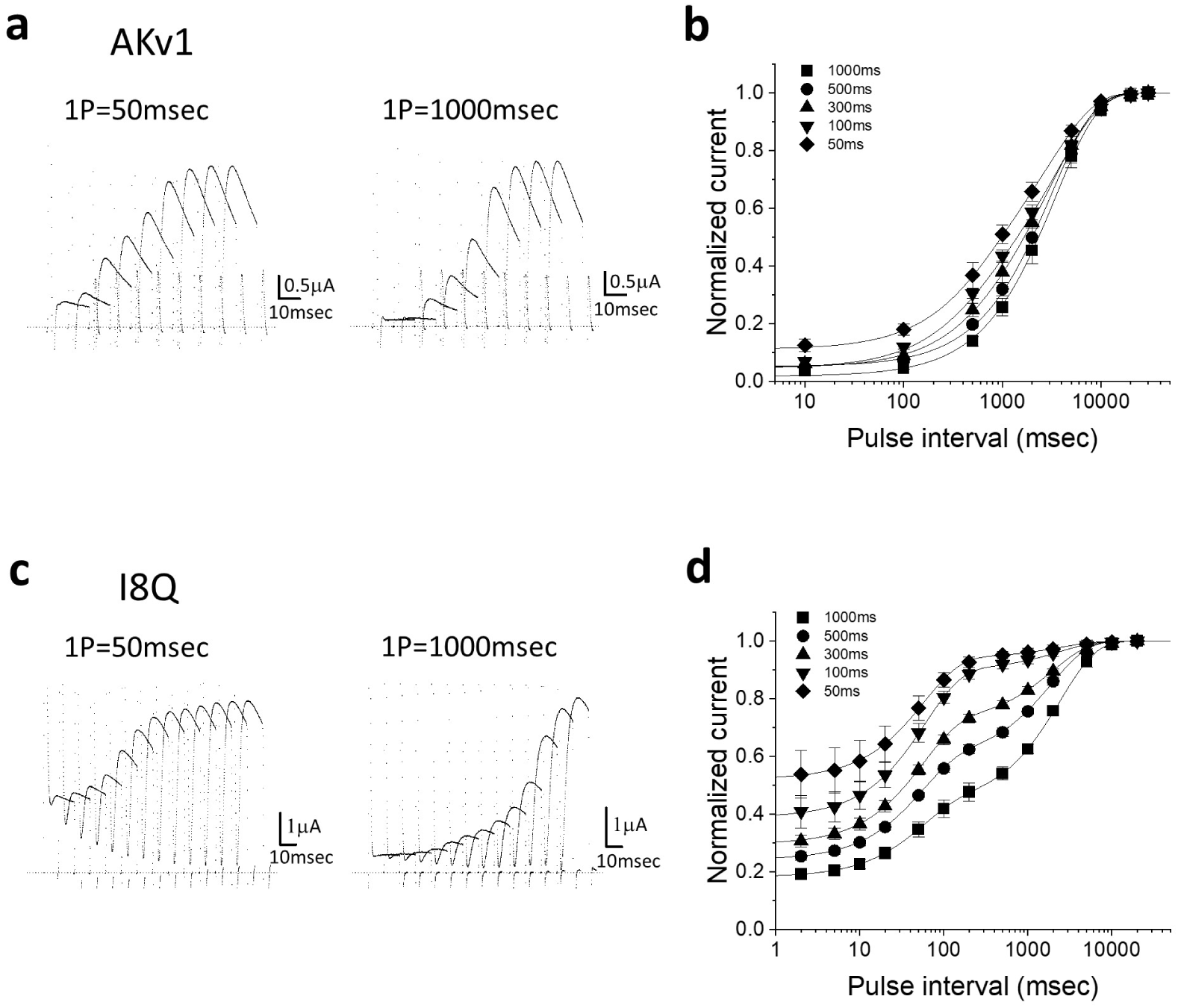
Comparison of the recovery from inactivation between AKv1 and I8Q. Conventional two pulse protocol was used to quantify the recovery. The prepulse (1P, +40 mV) was followed by the test pulse (2P, +40 mV, 20 msec) after the variable inter-pulse interval at -80 mV. The duration of 1P was 50, 100, 300, 500 or 1000 msec. **a:** Examples showing the recovery in AKv1. The currents during the 2P after the 1P of 50 or 1000 msec are superimposed with a constant time-shift for graphical reason. The actual inter-pulse intervals were as follows (msec): 10, 100, 500, 1000, 2000, 5000, 10000, 20000, 30000. **b:** The recovery time course of AKv1. Smooth lines are single exponential functions drawn by using mean parameters shown in Table 2. **c:** Examples showing the recovery in I8Q. The currents during the 2P are shown as in **a**. The actual inter-pulse interval were as follows (msec): 2, 5, 10, 20, 50, 100, 200, 500, 1000, 2000, 5000, 10000, 20000. **d:** The recovery time course of I8Q. In all cases, the recovery can be approximated by double exponential function. Smooth lines were double exponential functions drawn by using mean parameters shown in Table 2.

**Table 2.** Parameters for exponential fittings of the recovery from inactivation in AKv1 and I8Q.

| channel | pd | A1 | p | $\tau 1$ | p | A2 | p | $\tau 2$ | p | init | p | n |
| --- | --- | --- | --- | --- | --- | --- | --- | --- | --- | --- | --- | --- |
| AKv1 | 50 | -0.269±0.047 | - | 526.9±181.5 | - | -0.619±0.053 | - | 3249.2±461.6 | - | 0.116±0.019 | - | 5 |
| | 100 | -0.242±0.036 | 0.599 | 482.4±37.3 | 0.759 | -0.712±0.047 | 0.009 | 3714.3±528.3 | 0.391 | 0.061±0.012 | <0.001 | 5 |
| | 300 | -0.160±0.063 | 0.009 | 655.6±57.4 | 0.162 | -0.792±0.051 | <0.001 | 3418.4±427.0 | 0.950 | 0.051±0.021 | <0.001 | 5 |
| | 500 | - | - | - | - | -0.948±0.020 | <0.001 | 3161.8±473.2 | 0.996 | 0.052±0.020 | <0.001 | 4 |
| | 1000 | - | - | - | - | -0.993±0.023 | <0.001 | 3454.5±530.4 | 0.907 | 0.014±0.009 | <0.001 | 5 |
| I8Q | 50 | -0.430±0.074 | - | 57.7±2.5 | - | -0.062±0.016 | - | 2422.4±433.8 | - | 0.535±0.085 | - | 5 |
| | 100 | -0.522±0.037 | 0.008 | 59.6±6.7 | 0.853 | -0.105±0.018 | 0.009 | 2331.6±518.9 | >0.9 | 0.398±0.056 | 0.0004 | 5 |
| | 300 | -0.427±0.021 | >0.9 | 54.8±2.2 | 0.628 | -0.284±0.019 | <0.001 | 1958.1±174.3 | 0.159 | 0.303±0.021 | <1e-4 | 5 |
| | 500 | -0.354±0.032 | 0.042 | 52.9±3.0 | 0.223 | -0.409±0.018 | <0.001 | 1895.6±195.7 | 0.095 | 0.243±0.013 | <1e-4 | 6 |
| | 1000 | -0.254±0.031 | <0.01 | 53.4±3.4 | 0.301 | -0.565±0.040 | <0.001 | 2411.9±288.2 | >0.9 | 0.182±0.012 | <1e-4 | 5 |

Mean ± SD of parameters estimated by single or double exponential fitting of the recovery (see Materials and Methods) are shown.
pd: duration of the inactivating prepulse to +40 mV (msec)
A1, A2: the relative amplitudes of fast (A1) and slow (A2) exponential components
$\tau 1$ , $\tau 2$ : fast ( $\tau 1$ ) and slow ( $\tau 2$ ) time constants (msec)
init: the initial level of the normalized exponential recovery
p: p-values obtained by Dunnett's test. Parameters were compared between 50 msec pd and others.

It is well established that C-type inactivation of Kv channels is hindered by extracellular K^+^ [35, 40]. We therefor next compared the recovery from inactivation in ND96 and in high K^+^ solution (Fig. 3). We used a 1 sec prepulse to +40 mV in these experiments to quantify the recovery process. The recovery time course of AKv1 was well described by a single exponential in either condition and the recovery was much faster in high K^+^ condition (Fig. 3a, Table 3). The initial inactivated level after the prepulse (estimated by a minimum value in the exponential fitting, see ”init” in Table 3) in ND96 was slightly deeper than that in high K^+^ condition.

**Fig. 3.**
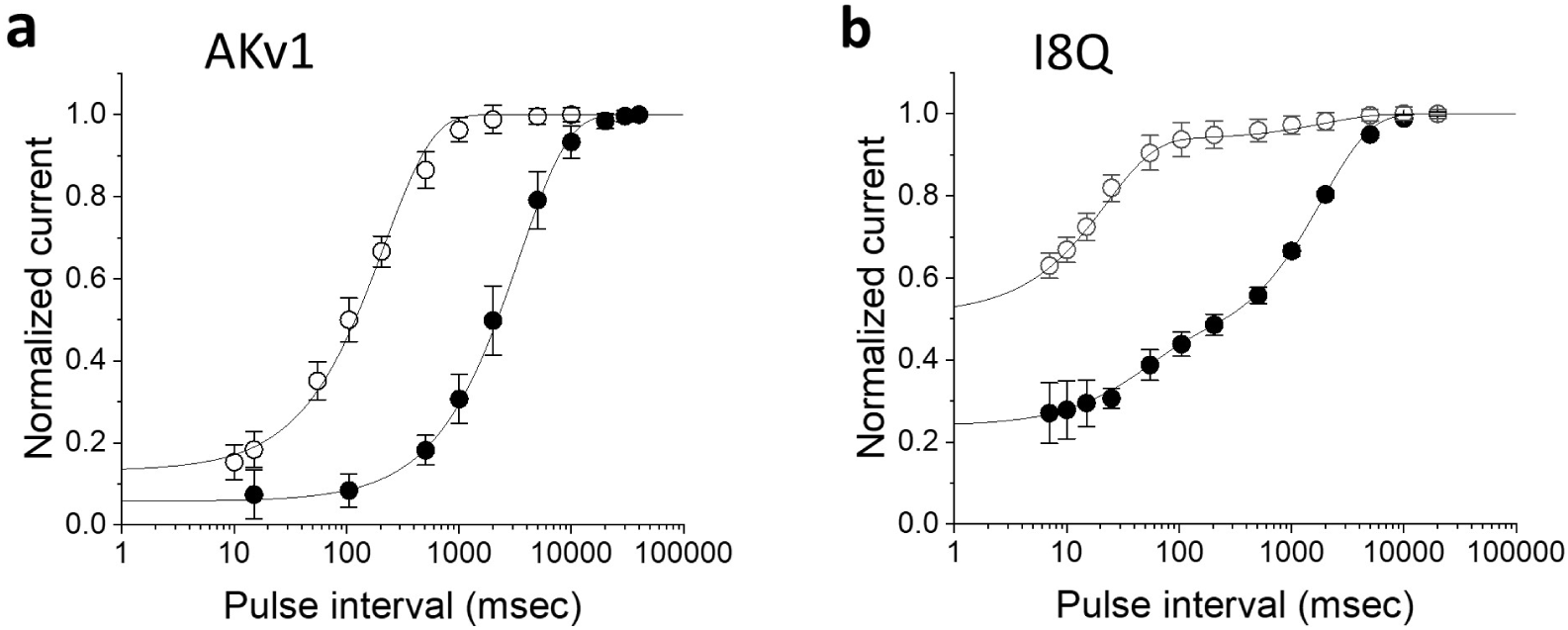
The recovery from inactivation of AKv1 (**a**) or I8Q (**b**) in ND96 and high K^+^ solution. The two pulse protocol with a 1sec prepulse (+40 mV) was used. The test pulse was +40 mV and the inter-pulse potential was -80 mV. Filled circle (ND96), Open circle (high K^+^). Smooth lines are exponential functions drawn by using mean parameters shown in Table 3.

**Table 3.**
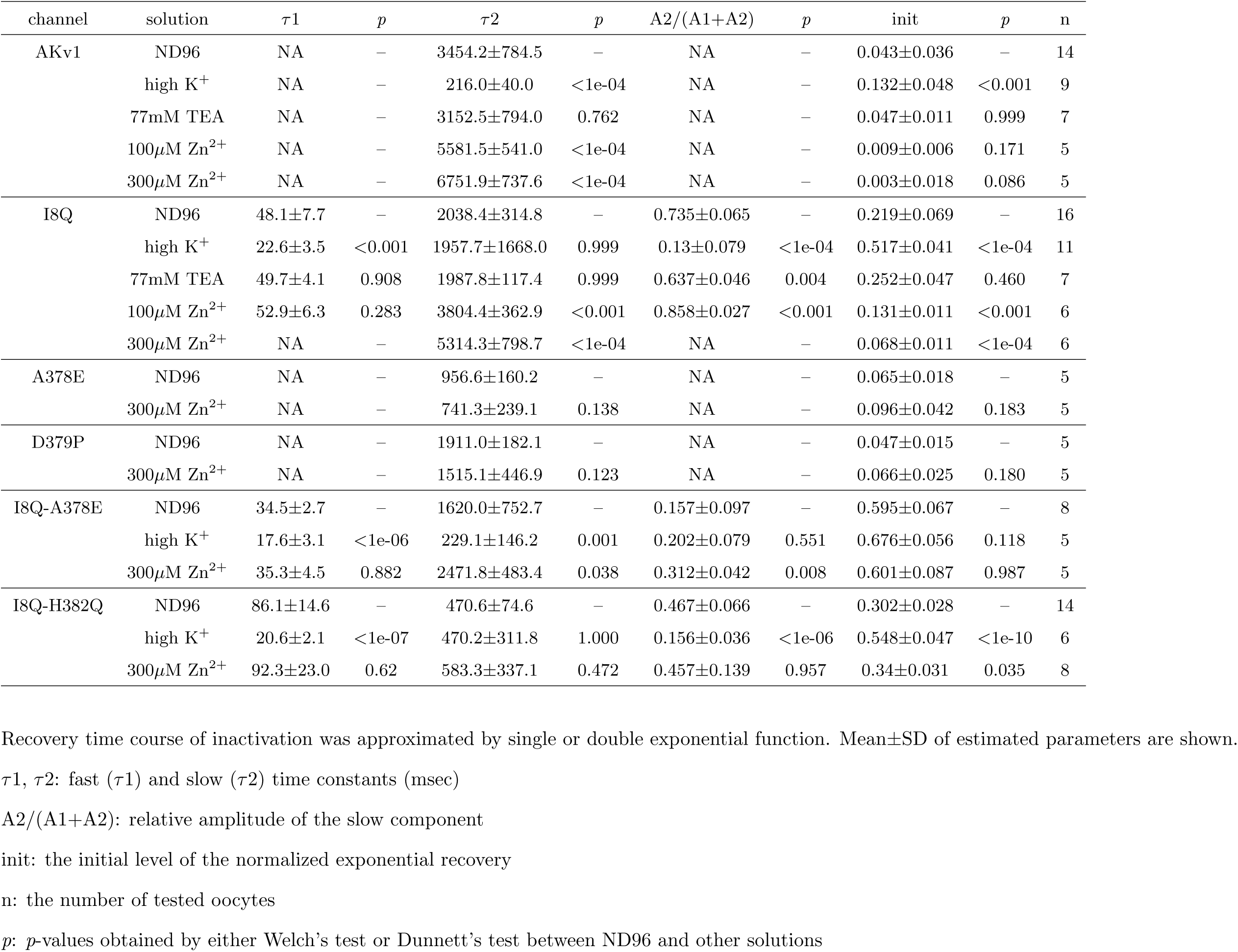
Comparison of the parameters for the recovery from inactivation in AKv1 and the I8Q mutants.

| channel | solution | $\tau_1$ | $p$ | $\tau_2$ | $p$ | A2/(A1+A2) | $p$ | init | $p$ | n |
| --- | --- | --- | --- | --- | --- | --- | --- | --- | --- | --- |
| AKv1 | ND96 | NA | - | 3454.2±784.5 | - | NA | - | 0.043±0.036 | - | 14 |
|  | high K<sup>+</sup> | NA | - | 216.0±40.0 | <1e-04 | NA | - | 0.132±0.048 | <0.001 | 9 |
|  | 77mM TEA | NA | - | 3152.5±794.0 | 0.762 | NA | - | 0.047±0.011 | 0.999 | 7 |
|  | 100μM Zn<sup>2+</sup> | NA | - | 5581.5±541.0 | <1e-04 | NA | - | 0.009±0.006 | 0.171 | 5 |
|  | 300μM Zn<sup>2+</sup> | NA | - | 6751.9±737.6 | <1e-04 | NA | - | 0.003±0.018 | 0.086 | 5 |
| I8Q | ND96 | 48.1±7.7 | - | 2038.4±314.8 | - | 0.735±0.065 | - | 0.219±0.069 | - | 16 |
|  | high K<sup>+</sup> | 22.6±3.5 | <0.001 | 1957.7±1668.0 | 0.999 | 0.13±0.079 | <1e-04 | 0.517±0.041 | <1e-04 | 11 |
|  | 77mM TEA | 49.7±4.1 | 0.908 | 1987.8±117.4 | 0.999 | 0.637±0.046 | 0.004 | 0.252±0.047 | 0.460 | 7 |
|  | 100μM Zn<sup>2+</sup> | 52.9±6.3 | 0.283 | 3804.4±362.9 | <0.001 | 0.858±0.027 | <0.001 | 0.131±0.011 | <0.001 | 6 |
|  | 300μM Zn<sup>2+</sup> | NA | - | 5314.3±798.7 | <1e-04 | NA | - | 0.068±0.011 | <1e-04 | 6 |
| A378E | ND96 | NA | - | 956.6±160.2 | - | NA | - | 0.065±0.018 | - | 5 |
|  | 300μM Zn<sup>2+</sup> | NA | - | 741.3±239.1 | 0.138 | NA | - | 0.096±0.042 | 0.183 | 5 |
| D379P | ND96 | NA | - | 1911.0±182.1 | - | NA | - | 0.047±0.015 | - | 5 |
|  | 300μM Zn<sup>2+</sup> | NA | - | 1515.1±446.9 | 0.123 | NA | - | 0.066±0.025 | 0.180 | 5 |
| I8Q-A378E | ND96 | 34.5±2.7 | - | 1620.0±752.7 | - | 0.157±0.097 | - | 0.595±0.067 | - | 8 |
|  | high K<sup>+</sup> | 17.6±3.1 | <1e-06 | 229.1±146.2 | 0.001 | 0.202±0.079 | 0.551 | 0.676±0.056 | 0.118 | 5 |
|  | 300μM Zn<sup>2+</sup> | 35.3±4.5 | 0.882 | 2471.8±483.4 | 0.038 | 0.312±0.042 | 0.008 | 0.601±0.087 | 0.987 | 5 |
| I8Q-H382Q | ND96 | 86.1±14.6 | - | 470.6±74.6 | - | 0.467±0.066 | - | 0.302±0.028 | - | 14 |
|  | high K<sup>+</sup> | 20.6±2.1 | <1e-07 | 470.2±311.8 | 1.000 | 0.156±0.036 | <1e-06 | 0.548±0.047 | <1e-10 | 6 |
|  | 300μM Zn<sup>2+</sup> | 92.3±23.0 | 0.62 | 583.3±337.1 | 0.472 | 0.457±0.139 | 0.957 | 0.34±0.031 | 0.035 | 8 |

Recovery time course of inactivation was approximated by single or double exponential function. Mean±SD of estimated parameters are shown.
$\tau_1$ , $\tau_2$ : fast ( $\tau_1$ ) and slow ( $\tau_2$ ) time constants (msec)
A2/(A1+A2): relative amplitude of the slow component
init: the initial level of the normalized exponential recovery
n: the number of tested oocytes
$p$ : $p$ -values obtained by either Welch's test or Dunnett's test between ND96 and other solutions

As shown in Fig. 2, the recovery time course of I8Q is approximated by double exponential function. The recovery from inactivation in high K^+^ was much faster than that in ND96 (Fig. 3b, Table 3). The fast *τ_rec_*became shorter and the recovery was dominated by the fast process (Table 3). Actually, a single exponential fitting may be good enough to approximate the recovery of I8Q in high K^+^ condition as described previously [51]. The recovery time course of I8Q in high K^+^ solution is rather close to the one obtained after the 50 msec prepulse in ND96, implying that both high K^+^ and a short prepulse prevent the channel entering into the more stable inactivated state.

Taken together, the results are consistent with previous suggestion that N-type inactivation of AKv1 is efficiently coupled to C-type inactivation [54]. On the other hand, the clear two components in the inactivation of I8Q in ND96 suggests that the N-C coupling in I8Q is less efficient.

### Comparison of the cumulative inactivation between AKv1 and I8Q

One of the interesting properties of AKv1 is a robust cumulative inactivation which can be observed by repetitive activation with short pulse (10–20 msec), during which the AKv1 current shows little inactivation [14]. A similar prominent cumulative inactivation is also observed in Kv1.4 [5]. Because the amino-terminal deletion mutant of AKv1 (ΔN) does not show the cumulative inactivation by such a short pulse, the cumulative inactivation of AKv1 depends on N-type inactivation [14]. To see the initial phase of the cumulative inactivation of AKv1, we compared the currents in response to two short depolarizing pulses (+40mV, 20msec) separated by variable inter-pulse interval at -40 to -160 mV (Fig. 4). When the inter-pulse potential was -40mV, the peak current at 2nd pulse became smaller initially by increasing the inter-pulse interval (Fig. 4a, black traces). By contrast, only the slow recovery was observed when the inter-pulse potential was -160 mV (Fig. 4a, gray traces). A counter-intuitive initial depression of the second current observed at the inter-pulse potential of -40 mV is not due to the development of steady state inactivation because AKv1 rarely inactivates by a sustained prepulse (ex., 2 sec) to -40 mV [15, 30, 52]. Similar phenomenon was seen in I8Q but much lesser extent (Fig. 4a). In Fig 4b and 4c, the relationships between the peak current ratio (I_2_*_nd_*/I_1_*_st_*) and the inter-pulse interval are illustrated at several inter-pulse potentials. The relationships were approximated by single or double exponential function (the parameters for exponential fittings are shown in Table S1). The reduction of the current at second pulse measured after the shortest interval (5 msec) as well as the estimated one by the fitting (1 msec after the 1st pulse) were *∼*40% in AKv1 and *∼*20% in I8Q, and the values were not meaningfully affected by the deeply hyperpolarized inter-pulse potential or high K^+^ condition, both of which can enhance the recovery from inactivation. Although a faster recovery in high K^+^ shown in Fig. 4 may reflect the reduced C-type inactivation [35, 40], the main cause of the development of inactivation during the inter-pulse interval in AKv1 is considered to be a voltage-independent N-type inactivation as suggested earlier [14]. In this scenario, I8Q seems to be less likely to enter and/or more easily get back from N-type inactivated state compared to AKv1.

**Fig. 4.**
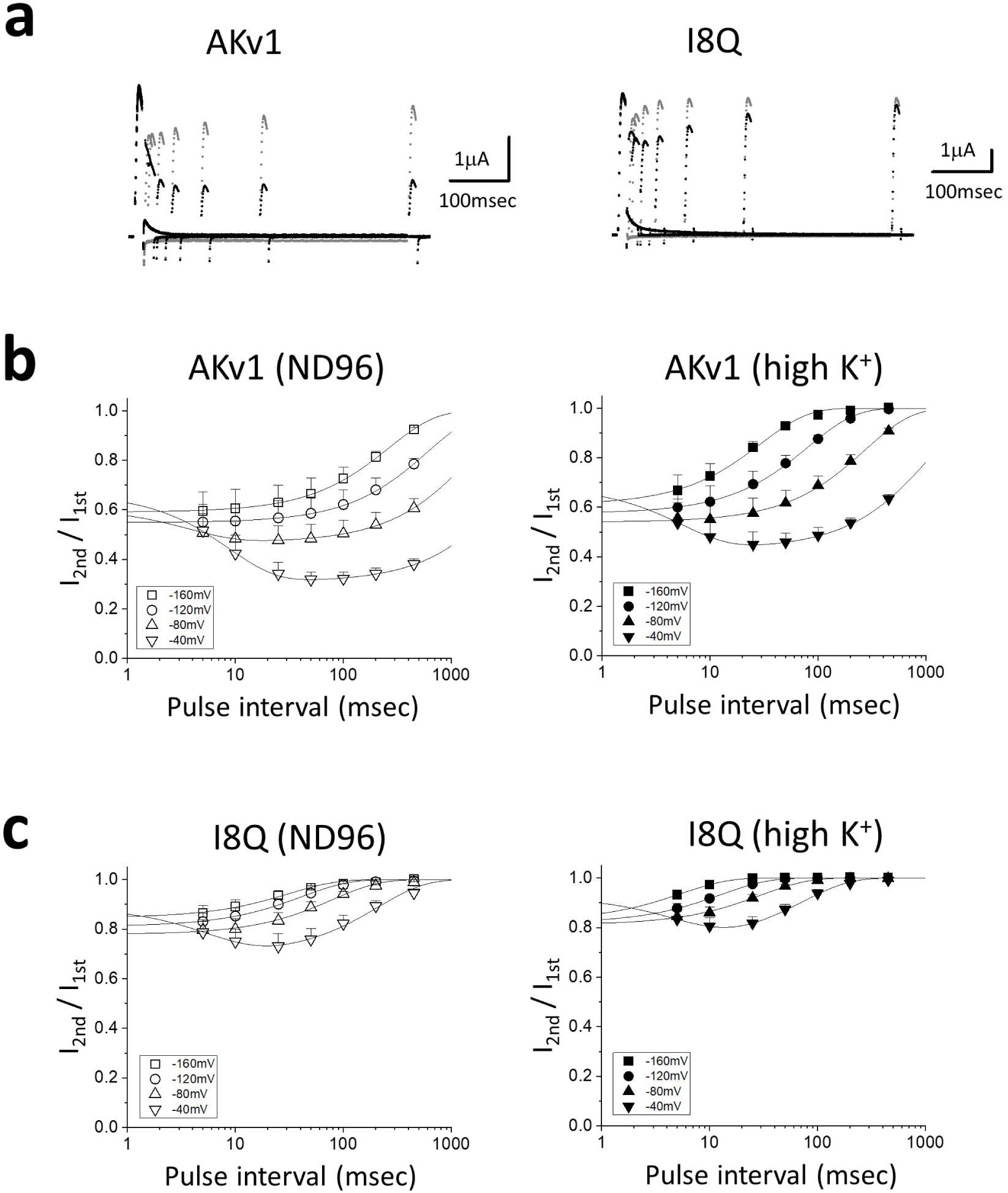
Cumulative inactivation assessed by two pulse protocol. Two short depolarizing pulses (+40mV, 20msec) were applied with a variable interval at the membrane potential of -40, -80, -120 or -160 mV. **a:** Examples in AKv1 and I8Q. The currents obtained by two identical pulses with an inter-pulse interval of 5, 10, 25 50 100, 200 or 500 msec are superimposed. The inter-pulse potential was -40 mV (black) or -160 mV (gray). **b:** The relationship between the ratio of two currents (I_2_*_nd_*/I_1_*_st_*) and the inter-pulse interval in AKv1. The data were approximated by either double exponential function (-40, -80 mV in ND96, -40 mV in high K^+^) or single exponential function (others) as described in Materials and Methods. Smooth lines are exponentials drawn by parameters shown in Table S1. **c:** The relationship between I_2_*_nd_*/I_1_*_st_* and the inter-pulse interval in I8Q. The data obtained by the inter-pulse potential of -40mV were fitted with double exponential function and others were fitted with single exponential function. Smooth lines are exponentials drawn by parameters shown in Table S1.

To obtain further insights, we next measured the gating currents of AKv1 and examined the recovery of gating charge after the short depolarizing pulse. To measure gating currents, we used W391F mutants of AKv1, I8Q and ΔN, which correspond to W434F mutant of *Drosophila Shaker* channel that had been extensively used to analyze the gating currents [50, 59, 69]. We measured the gating currents of AKv1 as well as its mutants by 10–40 msec depolarizing steps from the holding potential of -80 mV (see Materials and Methods). Fig. 5a and 5b shows examples of the families of gating currents of AKv1-W391F and I8Q-W391F and the charge-voltage relationships (Q-V curve) of the ON gating currents, respectively. Although the detailed analysis of the kinetics of the gating currents was not done in the present study, the waveforms of the gating currents of AKv1 and I8Q were basically similar and their charge-voltage relationships were close except for a slight depolarizing shift of the Q-V curve in I8Q as described in the voltage-dependent activation of I8Q. We examined the recovery of gating charges in AKv1-W391F, I8Q-W391F and ΔN-W391F by using two-pulse protocol similar to the one used to analyze the cumulative inactivation of ionic currents. For the gating current experiments, we used 10 msec depolarizing pulses to +40 mV separated by variable interval at -80 to -140 mV. The recovery time course was approximated by a single exponential function (the parameters for fittings are shown in Table S2). The recovery of the gating current of AKv1 became faster and more complete at larger hyperpolarized inter-pulse potential but some component of the ON gating charge did not seem to recover within *∼*100 msec (Fig. 5c). At the inter-pulse potential of -80 mV, *∼*40% of the ON gating charge seemed to be immobilized for *>*100 msec. Even at the inter-pulse potential of -120 mV or more, *∼*10% of the gating charge remained immobilized at least (Fig. 5c). The recovery of the gating charge was more complete in I8Q: the recovery of the charge following 100 msec interval was *∼*90% at -80 mV, *>*95% or more at -100 mV, and complete at -120 mV (Fig. 5d). In ΔN, the recovery at -80 mV was *>*95% and complete at -100 mV (Fig. 5e). On the other hand, the time constants for the charge recovery were almost identical among the channels: *∼*30–40 msec at -80mV, *∼*15–17 msec at -100mV, *∼*7–9 msec at -120mV, *∼*3–4 at -140mV (see Table S2).

**Fig. 5.**
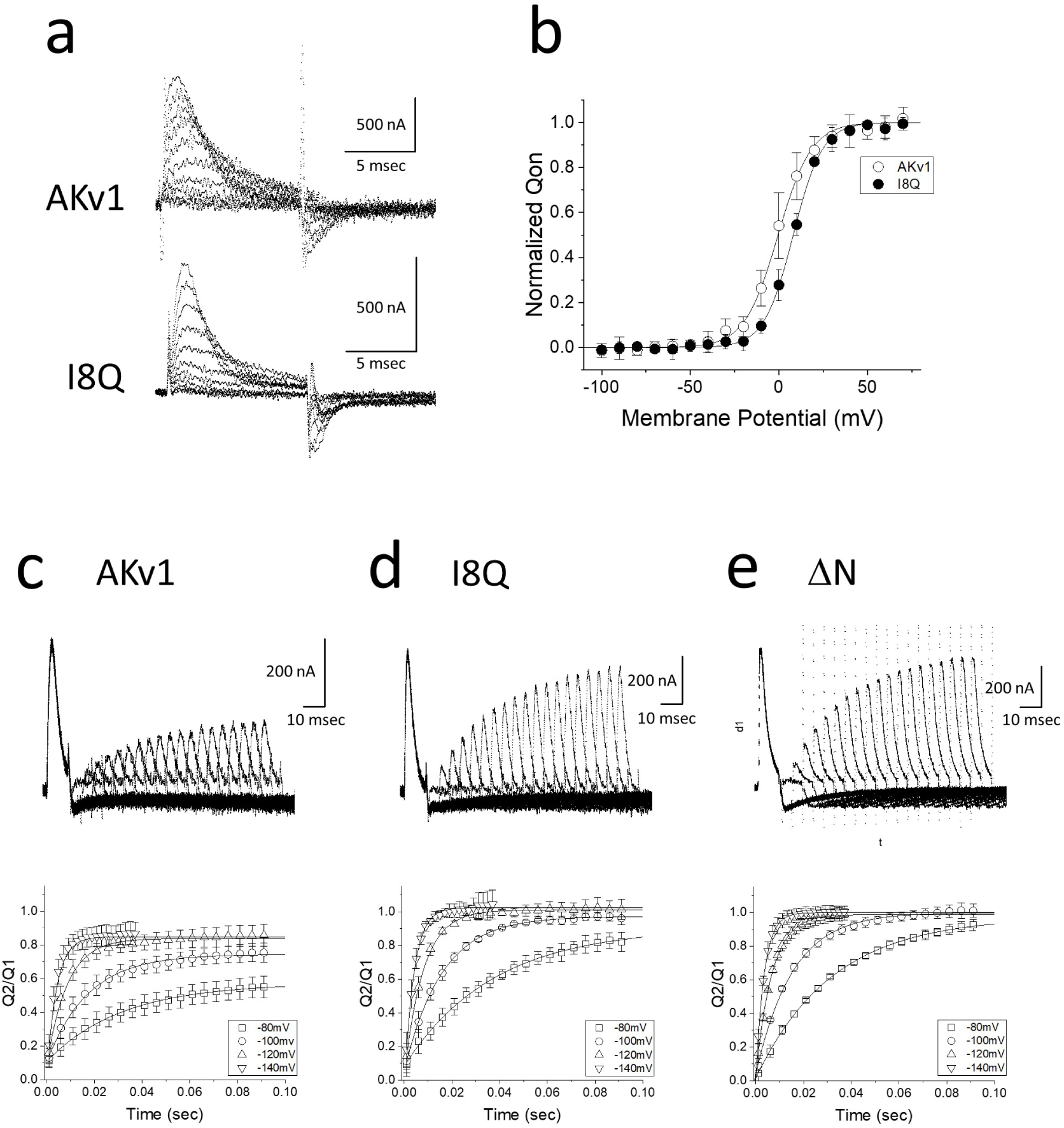
Comparison of the gating currents among AKv1, I8Q and ΔN. **a:** Examples of the family of gating currents in AKv1 and I8Q. 10 msec pulses (from -40 mV to 60 mV) were applied from the holding potential of -80 mV. Linear leak currents were subtracted by P/6 protocol as described in Materials and Methods. **b:** The normalized relationships between the ON gating charges and the membrane potentials. Smooth lines are fitted by Boltzmann equation as described in Materials and Methods. V*_half_* and k were as follows (mV): AKv1, -0.5*±*4.5, -9.4*±*1.8 (n=9); I8Q, 8.1*±*1.9, -8.3*±*1.1 (n=3). In **c** to **e**, the gating charge recovery of AKv1-W391F (**c**), I8Q-W391F (**d**) and ΔN-W391F (**e**) are shown. Two short depolarizing pulses (+40mV, 10msec) were applied with a variable interval at the membrane potential of -80, -100, -120 or -140mV. In each figure, upper panel shows the superimposed gating currents obtained by the inter-pulse potential of -80 mV. The currents obtained by two-pulse protocol with variable inter-pulse interval (5 msec increment from 1 msec) are superimposed. Lower panel shows the relationship between the ratio of the gating charges (Q2/Q1) and the inter-pulse interval. The gating charge was obtained by integrating the gating current during the depolarizing pulse. Smooth lines are exponentials drawn by mean parameters shown in Table S2.

The recovery time course of the ON gating charge can depend on the transitions among several closed states, transition to the opening, and the transitions after the opening including the inactivation. In the presence of 1 mM 4-AP which is known to arrest the opening transition of Kv channels [38, 43], the gating currents were apparently more symmetric and the gating charge recovery of AKv1 as well as ΔN became much faster (*τ* was *∼*2 msec at -80 mV, data not shown), suggesting that the recovery time courses observed in Fig. 5 are determined mostly by the transitions after the last closed state. An incomplete gating charge recovery observed in AKv1 can be explained by the gating charge immobilization following the inactivation [3, 6, 11, 68]. The fact that the charge immobilization following a 10 msec depolarizing pulse was prominent only in AKv1 but not in I8Q and ΔN is consistent with notions that the stability of N-type inactivation is a key for the cumulative inactivation of AKv1 by short repetitive pulses and that the N-type inactivated state of I8Q is not so stable as that of AKv1.

### Effects of TEA and Zn**^2+^** on the inactivation of AKv1

We next examined whether TEA and Zn^2+^ can modify the inactivation of AKv1. Although TEA is a well known blocker of some K^+^ channels [21] and also known to retard C-type inactivation of some Kv channels [7, 19], AKv1 is poorly sensitive to TEA [52] and even a high concentration of TEA (77mM) did not affect the inactivation induced by a 1 sec depolarizing pulse and its recovery in either AKv1 or I8Q (Table 1 and 3). On the other hand, Zn^2+^ appeared to affect C-type inactivation of AKv1 rather selectively as described below.

The peak current of AKv1 in response to a depolarizing step to +40 mV became slightly smaller in 100 or 300 *µ*M Zn^2+^ with a slight slowing of activation (Fig. 6a). The effect is qualitatively the same to that described in *Drosophila Shaker* channel and likely due to the shift of gating voltage range [17, 18, 61]. The current decay of AKv1 during a 1 sec depolarizing pulse was little affected by 100–300 *µ*M Zn^2+^ (Fig. 6a, Table 1). However, the recovery from inactivation of AKv1 in Zn^2+^ containing solution was slower than that in ND96 (Fig. 6b, Table 3), suggesting that the inactivated state becomes more stable in Zn^2+^ containing solution.

**Fig. 6.**
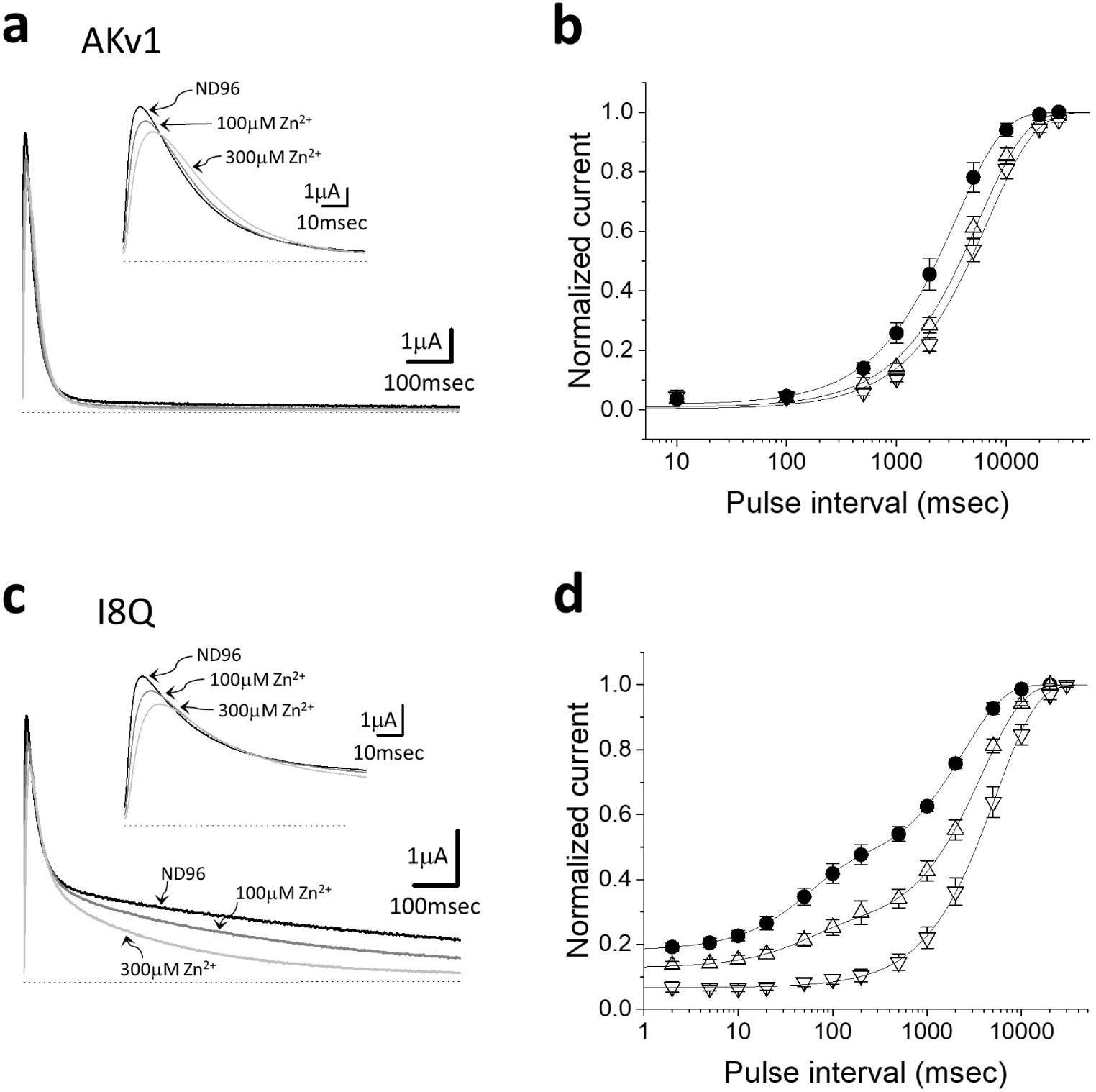
Effects of Zn^2+^ on the inactivation of AKv1 and I8Q. **a:** Effect of Zn^2+^ on the AKv1 current. The currents were evoked by 1 sec pulse to +40 mV from the holding potential of -80mV. The currents obtained in ND96, 100*µ*M Zn^2+^ and 300*µ*M Zn^2+^ are superimposed. The inset shows the time-expanded traces showing the initial 100 msec. Dotted lines indicate the zero current level. **b:** Effect of Zn^2+^ on the recovery from inactivation in AKv1. Filled circle (ND96), Up-pointing triangle (100*µ*M Zn^2+^), Down-pointing triangle (300*µ*M Zn^2+^). The two pulse protocol as shown in Fig. 3 was used. Smooth lines are single exponential functions drawn by using mean parameters shown in Table 3. **c:** Effect of Zn^2+^ on the I8Q current. The same pulse protocol as in **a**. The currents obtained in ND96, 100*µ*M Zn^2+^ and 300*µ*M Zn^2+^ are superimposed. The inset shows the time-expanded traces showing the initial 100 msec. Dotted lines indicate the zero current level. **d:** Effect of Zn^2+^ on the recovery from inactivation in I8Q. Filled circle (ND96), Up-pointing triangle (100*µ*M Zn^2+^), Down-pointing triangle (300*µ*M Zn^2+^). The two pulse protocol as shown in Fig. 3 was used. The recovery time course in ND96 or 100*µ*M Zn^2+^ was approximated by double exponential function and the recovery in 300*µ*M Zn^2+^ was approximated by single exponential. Smooth lines are drawn by using mean parameters shown in Table 3.

The peak current as well as the rate of rise of I8Q was also decreased slightly in Zn^2+^ containing solution (Fig. 6c). The fast component of inactivation of I8Q was rarely affected or marginally slower in Zn^2+^ containing solution (Fig. 6c, Table 1). By contrast, the slow component of inactivation was markedly accelerated by Zn^2+^ (Fig. 6c, Table 1). The recovery from inactivation of I8Q was inhibited in Zn^2+^ containing solution (Fig. 6d). The fast *τ_rec_*of I8Q in 100 *µ*M Zn^2+^ was similar to that in ND96, but the slow *τ_rec_*was much slower and the relative amplitude of the slow component became larger (Table 3). In 300 *µ*M Zn^2+^, the recovery time course of I8Q was well approximated by a single exponential with a slow time constant which was close to *τ_rec_* of AKv1 in Zn^2+^ containing solution (see Table 3). These results indicate that the slow inactivation of I8Q is specifically augmented by external Zn^2+^, resulting in the acceleration of the N-C coupling in I8Q.

### Effects of high K**^+^**, TEA, and Zn**^2+^** on the inactivation of amino-terminal deletion mutant of AKv1

We next used the amino-terminal deletion mutant of AKv1 (ΔN) to examine the C-type inactivation of AKv1 in the absence of N-type inactivation. Because the inactivation of ΔN was quite slow, we used 10 sec pulse to +40 mV to analyze the inactivation of ΔN. As shown in Fig. 7, ΔN shows substantial inactivation during 10 sec pulse to +40 mV albeit not complete. We first checked the effect of high K^+^ solution and TEA on the inactivation. As seen in Fig. 7a, the ΔN current in high K^+^ solution was smaller than that in ND96 as expected from the reduced driving force. The current decay in high K^+^ solution was slower and less complete as seen more clearly in the normalized current trace (a broken line in Fig. 7a, Table 4). Although it was possible to fit the inactivation of ΔN by a single exponential function in some experiments, the inactivation of ΔN was usually better approximated by double exponential function. However, because the slow component was dominant in the inactivation of ΔN as well as its mutants (usually *>*90%, see Table 4), we call the slow time constant *τ_inacti_*. *τ_inacti_*of ΔN in high K^+^ condition was much larger than that in ND96, which is a hallmark of C-type inactivation [35, 40].

**Fig. 7.**
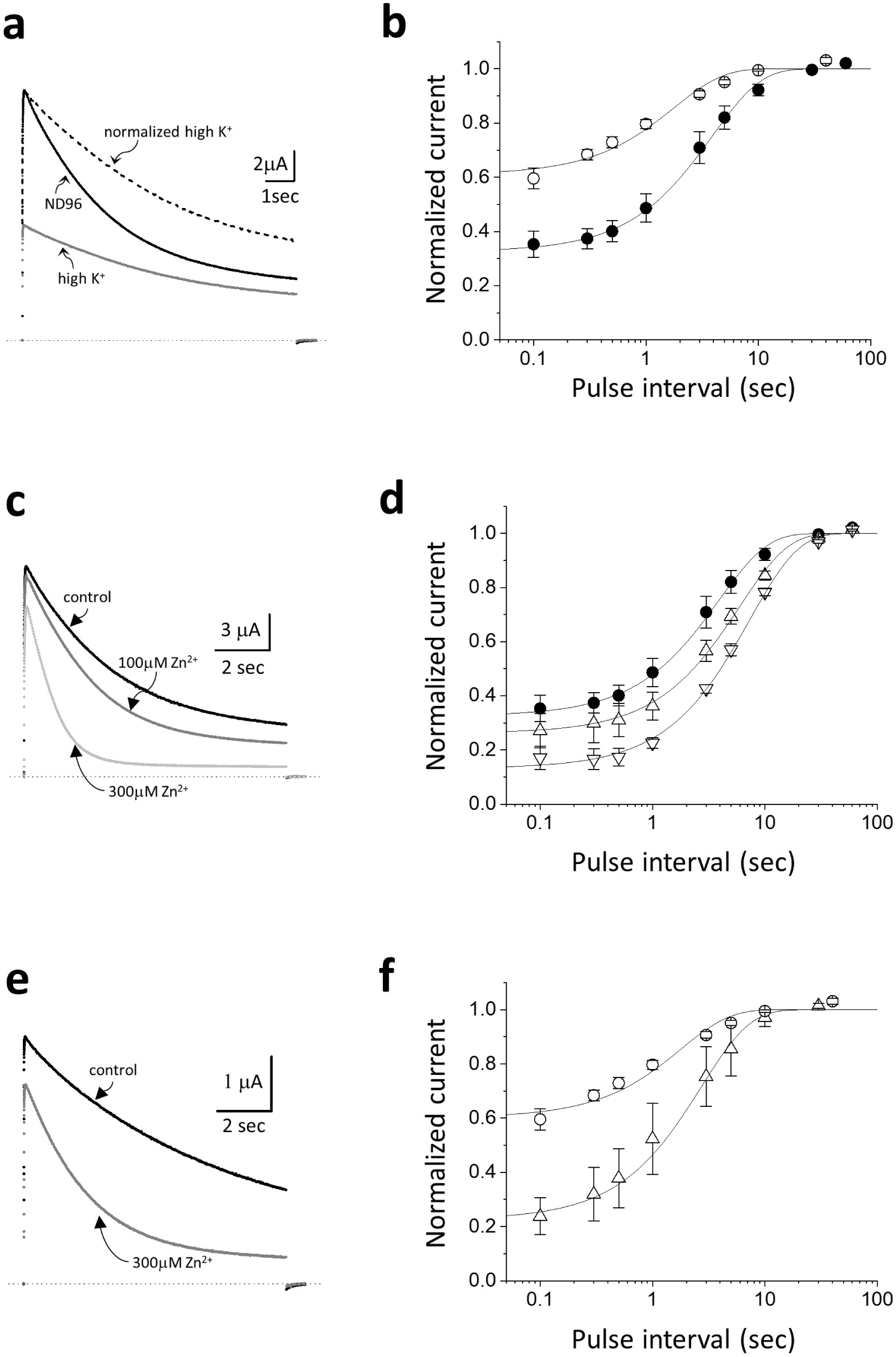
Effects of high K^+^ solution and Zn^2+^ on the inactivation of the amino-terminal deletion mutant of AKv1 (ΔN). **a:** Effect of high K^+^ solution on the ΔN current. The current traces shown here and in **c** and **e** were evoked by 10 sec pulse to +40 mV from the holding potential of -80mV. Dotted lines in this and in **c** and **e** indicate the zero current level. The current trace shown by broken line is a normalized trace of the current in high K^+^. **b:** Effect of high K^+^ solution on the recovery from inactivation in ΔN. Filled circle (ND96), Open circle (high K^+^). A 10 sec prepulse to +40 mV was followed by the test pulse to +40 mV after the variable inter-pulse interval at -80 mV. Smooth lines in this and in **d** and **f** are single exponentials drawn by using mean parameters shown in Table 5. **c:** Effect of Zn^2+^ on the ΔN current in ND96. **d:** Effect of Zn^2+^ on the recovery from inactivation in ND96. Filled circle (ND96, the same data shown in **b**), Up-pointing triangle (100*µ*M Zn^2+^), Down-pointing triangle (300*µ*M Zn^2+^). **e:** Effect of 300 *µ*M Zn^2+^ on the ΔN current in high K^+^ solution. **f:** Effect of 300 *µ*M Zn^2+^ on the recovery from inactivation in high K^+^ solution. Circle (high K^+^, the same data shown in **b**), Triangle (300*µ*M Zn^2+^ in high K^+^).

**Table 4.**
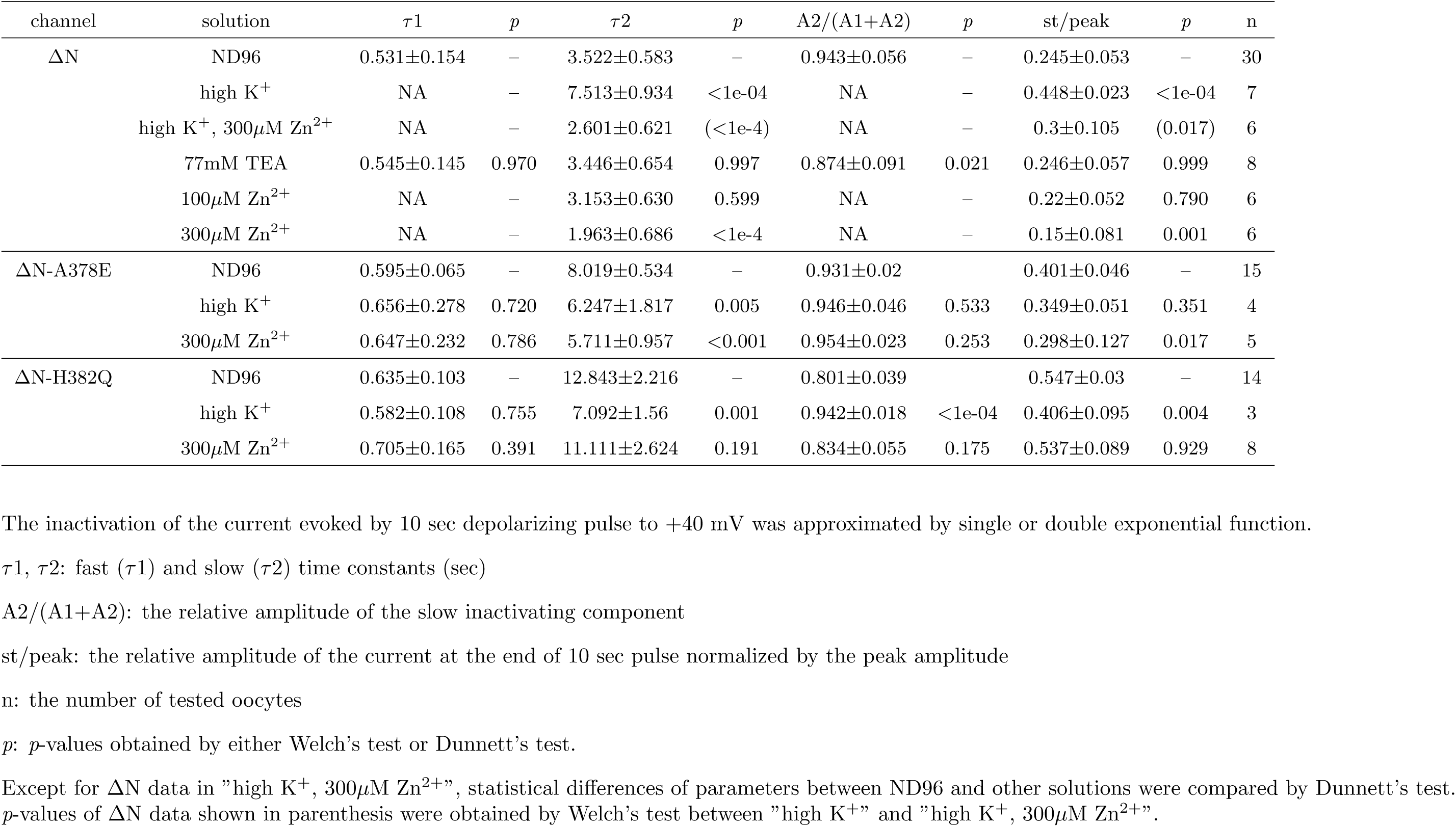
Inactivation of ΔN and the ΔN mutants.

**Table 5.** Comparison of the parameters for the recovery from inactivation in ΔN and ΔN mutants.

| channel | solution | $\tau 1$ | $p$ | $\tau 2$ | $p$ | A2/(A1+A2) | $p$ | init | $p$ | n |
| --- | --- | --- | --- | --- | --- | --- | --- | --- | --- | --- |
| $\Delta N$ | ND96 | NA | - | 3.491±0.968 | - | NA | NA | 0.325±0.038 | - | 11 |
| | high K<sup>+</sup> | NA | - | 1.781±0.175 | 0.002 | NA | NA | 0.61±0.03 | <0.001 | 5 |
| | high K<sup>+</sup>, 300μM Zn<sup>2+</sup> | NA | - | 2.716±1.152 | (0.203) | NA | NA | 0.226±0.07 | (<0.001) | 4 |
| | 77mM TEA | NA | - | 3.368±0.911 | 0.996 | NA | NA | 0.37±0.045 | 0.245 | 5 |
| | 100μM Zn<sup>2+</sup> | NA | - | 5.94±0.566 | <1e-04 | NA | NA | 0.262±0.068 | 0.054 | 5 |
| | 300μM Zn<sup>2+</sup> | NA | - | 7.334±0.619 | <1e-04 | NA | NA | 0.132±0.038 | <0.001 | 3 |
| $\Delta N$-A378E | ND96 | 0.195±0.025 | - | 9.142±1.021 | - | 0.535±0.095 | - | 0.556±0.032 | - | 8 |
| | high K<sup>+</sup> | 0.173±0.026 | 0.963 | 8.176±0.28 | 0.341 | 0.501±0.06 | 0.712 | 0.343±0.074 | <1e-04 | 4 |
| | 300μM Zn<sup>2+</sup> | 0.358±0.28 | 0.144 | 8.101±1.717 | 0.249 | 0.618±0.048 | 0.146 | 0.531±0.061 | 0.641 | 5 |
| $\Delta N$-H382Q | ND96 | 0.23±0.06 | - | 9.849±1.847 | - | 0.476±0.048 | - | 0.565±0.05 | - | 9 |
| | high K<sup>+</sup> | 0.169±0.036 | 0.263 | 7.176±0.626 | 0.138 | 0.494±0.061 | 0.905 | 0.441±0.028 | 0.002 | 3 |
| | 300μM Zn<sup>2+</sup> | 0.16±0.068 | 0.079 | 6.188±2.755 | 0.009 | 0.382±0.099 | 0.045 | 0.592±0.046 | 0.470 | 6 |

Recovery time course was approximated by single or double exponential function. Mean ± SD of estimated parameters are shown.
$\tau 1$ , $\tau 2$ : fast ( $\tau 1$ ) and slow ( $\tau 2$ ) time constants (msec)
A2/(A1+A2): relative amplitude of the slow component
init: the initial level of the normalized exponential recovery
n: the number of tested oocytes
$p$ : $p$ -values obtained by either Welch's test or Dunnett's test.
Except for $\Delta N$ data in "high K<sup>+</sup>, 300 $\mu$ M Zn<sup>2+</sup>", statistical differences of parameters between ND96 and other solutions were compared by Dunnett's test. $p$ -values of $\Delta N$ data shown in parenthesis were obtained by Welch's test between "high K<sup>+</sup>" and "high K<sup>+</sup>, 300 $\mu$ M Zn<sup>2+</sup>".

The recovery from inactivation examined by two pulse protocol (the inactivating pre-pulse was 10 sec to +40 mV) is shown in Fig. 7b. In ND96, the recovery time course was approximated by a single exponential with *τ_rec_*of *∼*4 sec (Table 5). The *τ_rec_*of ΔN was close to that of AKv1 in ND96 (see Fig. 3, Table 3), consistent with a notion that the recovery time course of AKv1 is normally determined by the recovery from C-type inactivated state in low K^+^ condition [54]. The *τ_rec_* of ΔN in high K^+^ solution became *∼*2 sec which was much shorter than that in ND96 (Table 5). The *τ_rec_* of ΔN in high K^+^ was, however, much longer than that of AKv1 in high K^+^ (*∼*200 msec), suggesting that the recovery of AKv1 in high K^+^ is not from C-type inactivated state but likely from N-type inactivated state [14, 54].

We next checked the effect of TEA on the C-type inactivation of ΔN. Although TEA is well known to inhibit C-type inactivation in some Kv channels [7, 19], external TEA had minor effect, if any, on the inactivation of ΔN as well as its recovery from inactivation (see Table 4, 5). By contrast, external Zn^2+^ markedly accelerated the inactivation of ΔN (Fig. 7c). Zn^2+^ depressed the peak current slightly, which is likely due to both the gating shift by Zn^2+^ as described above and the accelerated inactivation. In the presence of Zn^2+^, the current decay of ΔN during the 10 sec pulse was faster and the inactivation was more complete (Fig. 7c, Table 4). As expected from the more complete inactivation, the recovery from inactivation became slower in Zn^2+^ containing solution (Fig. 7d, Table 5). To address whether high K^+^ inhibits the action of Zn^2+^, we next examined the effect of Zn^2+^ on the inactivation of ΔN in high K^+^ condition. Even in high K^+^ condition, 300 *µ*M Zn^2+^ markedly accelerated the inactivation of ΔN (Fig. 7e, Table 4) and inhibited the recovery from inactivation (Fig 7f, Table 5), indicating that the effect of Zn^2+^ is not blocked by external K^+^.

### Effects of high K**^+^** and Zn**^2+^** on the inactivation of A378E mutants

If the effect of external Zn^2+^ on the inactivation of AKv1 is mostly dependent on C-type inactivation, Zn^2+^ should have little effect on the inactivation of the mutant channels which have impaired C-type mechanism. We previously examined several mutants around the pore turret region of AKv1, and found that some mutants show deteriorated C-type inactivation with intact N-type mechanism [60]. Among the previously described mutants, we checked two mutants, A378E and D379P, to see the effect of Zn^2+^ on their inactivation.

The *τ_inacti_*of A378E or D379P in response to a 1 sec depolarizing pulse was similar to that of AKv1 [60] but the *τ_rec_* was faster (Table 3), implying that C-type inactivation of these turret mutants is less stable compared to AKv1. We examined whether Zn^2+^ can affect the inactivation of A378E and D379P, and found that 300 *µ*M Zn^2+^ does not affect the inactivation of A378E and D379P as well as their recovery at all (Table 3).

To see the effect of Zn^2+^ on C-type inactivation more directly, we examined an aminoterminal deletion mutant of A378E (ΔN-A378E). The inactivation of ΔN-A378E was slower and less complete compared to ΔN (Fig. 8a, Table 4), showing the defectiveness of C-type mechanism. As in ΔN, the inactivation of ΔN-A378E was approximated by two exponential function and the slow component having *τ_inacti_* of *∼*8 sec was dominant (Table 4). Unexpectedly, however, the inactivation of ΔN-A378E became faster in high K^+^ solution (see a normalized trace in Fig. 8a, Table 4), suggesting that the slow inactivation of ΔN-A378E is somewhat different from the C-type inactivation of ΔN (see Discussion). Although Zn^2+^ slightly shortened *τ_inacti_*of ΔN-A378E (Table 4), the overall inactivation of ΔN-A378E was not seemed to be so different (Fig. 8b). The recovery from inactivation in ΔN-A378E was approximated by double exponential function: the fast and the slow *τ_rec_* were *∼*200 msec and *∼*10 sec, respectively (Table 5). Because the slow component accounted for only *∼*50% of the recovery in ΔN-A378E, the time required for full recovery after a 10 sec depolarizing prepulse was similar to that in ΔN. Both the fast and slow components in the recovery were not affected meaningfully in either high K^+^ solution or 300 *µ*M Zn^2+^ except for the initial inactivated level after the 10 sec prepulse in high K^+^ (see Fig. 8a, 8c, and Table 5). These results suggest that the A378 mutation modifies C-type mechanism of ΔN, and that the inactivation becomes much less sensitive to Zn^2+^. To better see the effect of A378E mutation on the N-C coupling, we introduced A378E mutation in I8Q. The inactivation of I8Q-A378E was approximated by two exponential function as in I8Q but it was much less complete. In high K^+^ solution, the inactivation of I8Q-A378E was inhibited further (Fig. 8d). In ND96, the fast *τ_inacti_*of I8Q-A378E was similar to I8Q as well as AKv1 but the slow *τ_inacti_*was much longer than that of I8Q (Table 1) and the relative amplitude of the current at the end of 1 sec pulse to +40 mV was more than 0.5 (see ”st/peak” in Table 1). By contrast to ΔN-A387E, these results are consistent with a notion that the slow component of inactivation in I8Q-A378E still depends on C-type mechanism.

**Fig. 8.**
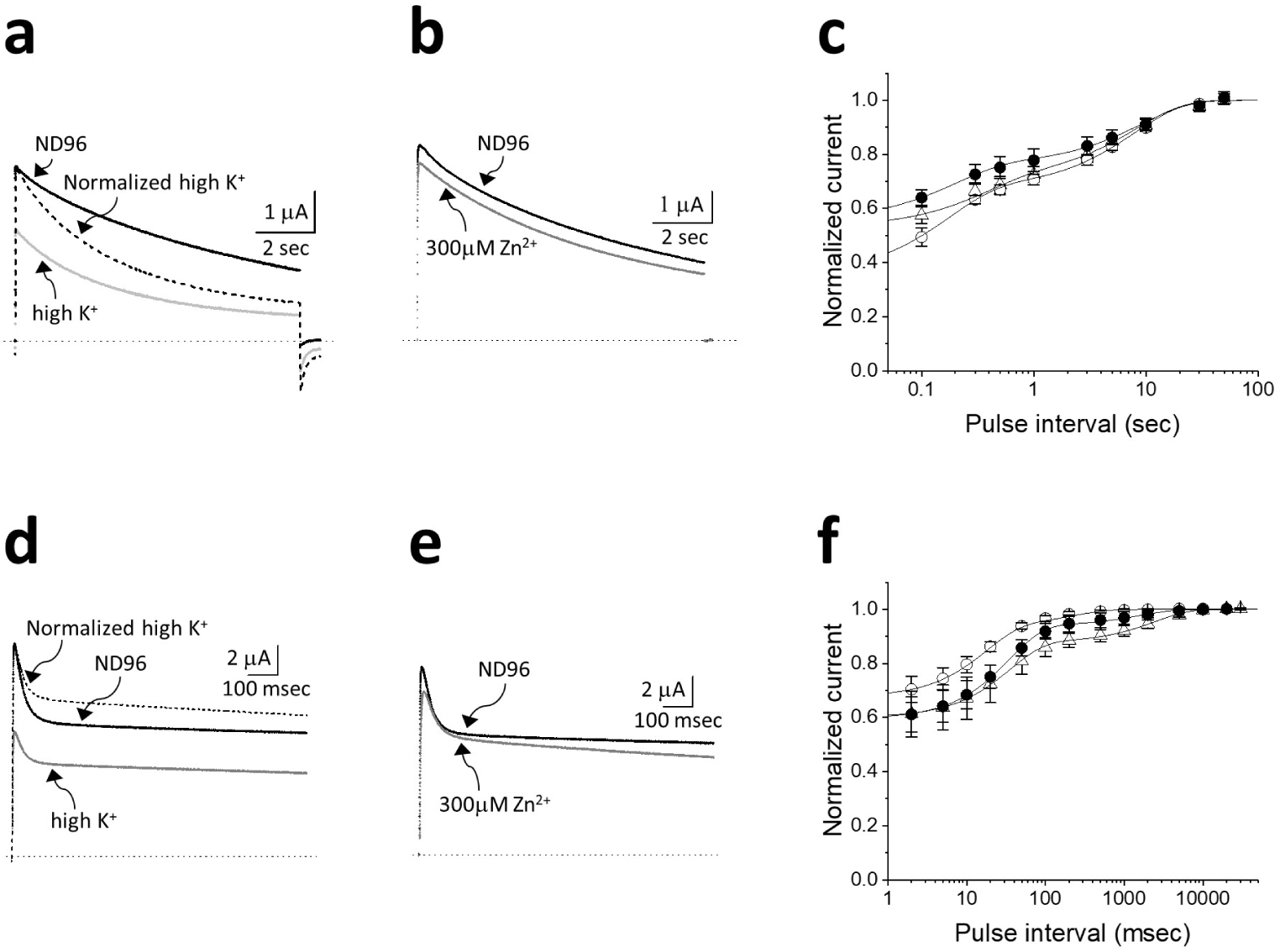
Effects of high K^+^ solution and Zn^2+^ on the inactivation of the A378E mutants. **a:** Effect of high K^+^ solution on the ΔN-A378E current. The currents were evoked by 10 sec pulse to +40 mV from the holding potential of -80mV. Dotted line indicates the zero current level. The current trace shown by broken line is a normalized trace of the current in high K^+^. **b:** Effect of 300 *µ*M Zn^2+^ on the ΔN-A378E current. **c:** Effect of high K^+^ solution and Zn^2+^ on the recovery from inactivation in ΔN-A378E. Filled circle (ND96), Open circle (high K^+^), Open triangle (300*µ*M Zn^2+^). The protocol was as describe in the legend of Fig. 7b. Smooth lines are double exponentials drawn by using mean parameters shown in Table 5. **d:** Effect of high K^+^ solution on the I8Q-A378E current. The currents were evoked by 1 sec pulse to +40 mV from the holding potential of -80mV. Dotted line indicates the zero current level. The current trace shown by broken line is a normalized trace of the current in high K^+^. **e:** Effect of 300 *µ*M Zn^2+^ on the I8Q-A378E current. **f:** Effects of high K^+^ and Zn^2+^ on the recovery from inactivation in I8Q-A378E. The protocol was similar to the one described in Fig. 3. Filled circle (ND96), Open circle (high K^+^), Up-pointing triangle (300 *µ*M Zn^2+^). Smooth lines are double exponentials drawn by using mean parameters shown in Table 3.

As in I8Q, the recovery from inactivation of I8Q-A378E was fitted by double exponential function, but the time course of recovery was much faster because the most recovery of this mutant was from a fast process (Fig. 8f, Table 3). In high K^+^ solution, the recovery of I8Q-A378E was further sped up with shorter *τ_rec_*s (Table 3). 300 *µ*M Zn^2+^ affected the inactivation of I8Q-A378E but the effect was diminished compared to the effect of Zn^2+^ on the inactivation of I8Q (Fig. 8e, Table 1). Zn^2+^ slightly inhibited the recovery from inactivation by enhancing the slow component but the effect was modest (Fig. 8f, Table 3). Collectively, the slow inactivation of I8Q is hindered by the A378E mutation but its sensitivities to K^+^ and Zn^2+^ still remain in I8Q-A378E.

### Effects of high K**^+^** and Zn**^2+^** on the inactivation of **Δ**N-H382Q and I8Q-H382Q

In Kv1.5, Zn^2+^ is supposed to bind to a histidine residue in the pore turret (H463 in hKv1.5) and stabilize the C-type inactivated state [12, 31]. Because AKv1 also has a histidine at homologous site (H382), we next examined H382Q mutants to see whether its inactivation is modified by external K^+^ and Zn^2+^. In ND96, the inactivation of ΔN- H382Q was extremely slow compared to ΔN or ΔN-A378E (Fig. 9a, Table 4). The slow inactivation of ΔN-H382Q became faster in high K^+^ solution as observed in ΔN-A378E (see a normalized trace in Fig. 9a, Table 4), suggesting similar functional modification of C-type inactivation. On the other hand, external Zn^2+^ rarely affected the inactivation of ΔN-H382Q (Fig. 9b, Table 4). Fig. 9c compares the recovery from inactivation of ΔN- H382Q in ND96, high K^+^ and 300 *µ*M Zn^2+^. The recovery time course was approximated by double exponential function and the parameters are shown in Table 5. In either high K^+^ solution or 300 *µ*M Zn^2+^, the recovery from inactivation was rather similar to that in ND96 except for some minor differences in *τ_rec_* or the initial inactivated level (Table 5). Overall, the inactivation of ΔN-H382Q was more sluggish compared to ΔN and the recovery from inactivation was less affected by high K^+^ and Zn^2+^.

**Fig. 9.**
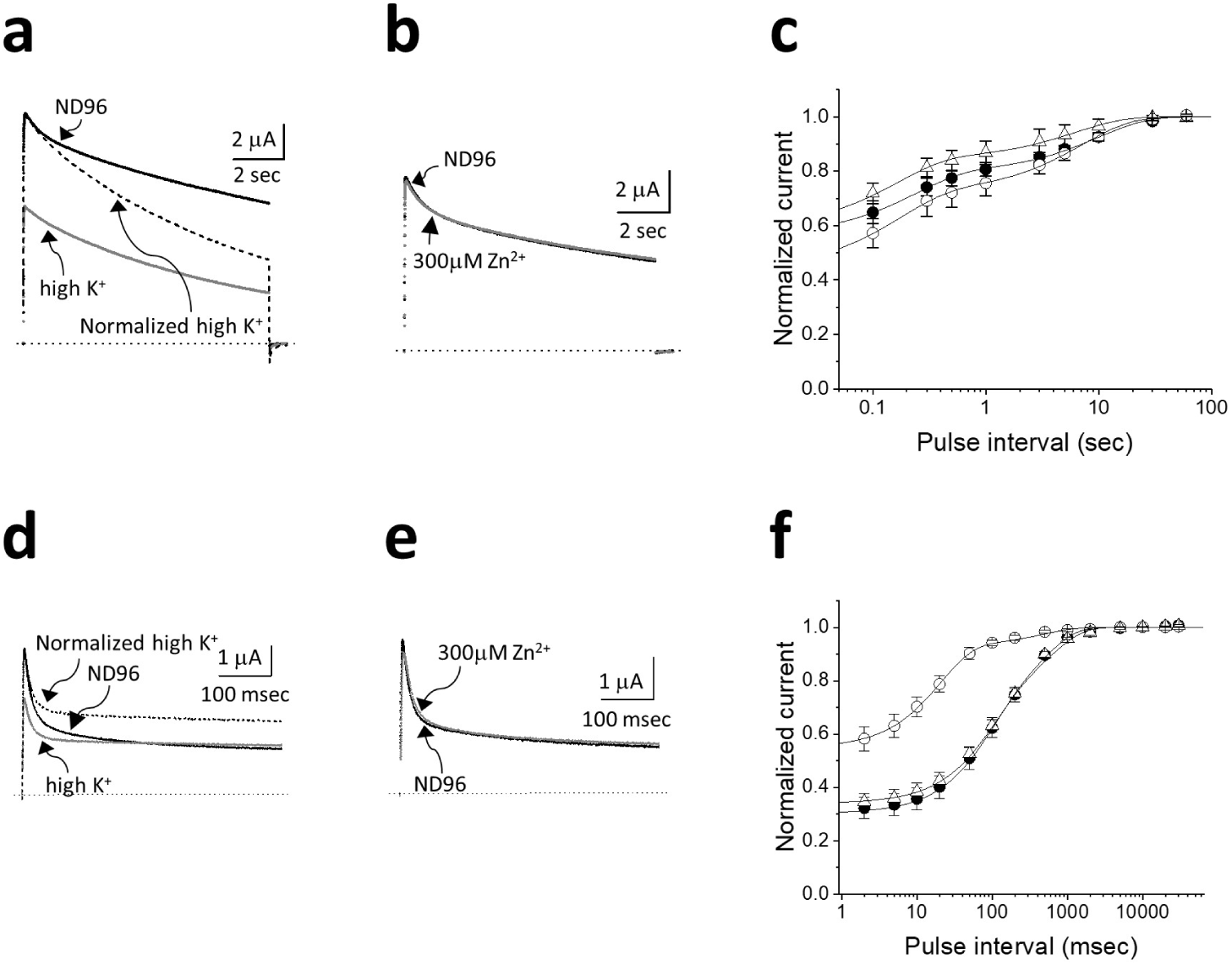
The inactivation of the H382Q mutants. **a:** Effect of high K^+^ solution on the ΔN- H382Q current. The currents were evoked by 10 sec pulse to +40 mV from the holding potential of -80mV. Dotted line indicates the zero current level. The current trace shown by broken line is a normalized trace of the current in high K^+^. **b:** Effect of 300 *µ*M Zn^2+^ on the ΔN-H382Q current. **c:** Effect of high K^+^ solution and Zn^2+^ on the recovery from inactivation in ΔN-H382Q. Filled circle (ND96), Open circle (high K^+^), Open triangle (300*µ*M Zn^2+^). The protocol was as describe in Fig. 7b. The recovery time course was approximated by double exponential function. Smooth lines are exponentials drawn by using mean parameters shown in Table 5. **d:** Effect of high K^+^ solution on the I8Q-H382Q current. The currents were evoked by 1 sec pulse to +40 mV from the holding potential of -80mV. Dotted line indicates the zero current level. The current trace shown by broken line is a normalized trace of the current in high K^+^. **e:** Effect of 300 *µ*M Zn^2+^ on the I8Q-H382Q current. **f:** Effects of high K^+^ and Zn^2+^ on the recovery from inactivation in I8Q-H382Q. The protocol was similar to the one described in Fig. 3. Filled circle (ND96), Open circle (High K^+^), Open triangle (300 *µ*M Zn^2+^). Smooth lines are double exponentials drawn by using mean parameters shown in Table 3.

We next made a mutant, I8Q-H382Q and examined its inactivation. The inactivation of I8Q-H382Q showed two components as in I8Q. The fast *τ_inacti_*of I8Q-H382Q was similar to that of I8Q (Fig. 9d, Table 1), but the slow *τ_inacti_* was faster and the total inactivating component during 1 sec pulse seemed to be slightly smaller than I8Q (see Table 1). The inactivation of I8Q-H382Q was still better approximated by double exponential function in high K^+^ solution, but the slow *τ_inacti_* became very long and the slow component was diminished (see Table 1), indicating that the slow inactivation of I8Q-H382Q is hindered by high K^+^ as in I8Q. Although two exponential function was still required to better fit the recovery time course of the inactivation of I8Q-H382Q in ND96 (Fig. 9f), the slow *τ_rec_* was much shorter than that of I8Q and the amplitude of slow component was smaller (Table 3). In high K^+^ condition, the recovery of I8Q-H382Q was dominated by a fast process having a time constant of *∼*20 msec and the overall recovery time course was essentially the same to that of I8Q in high K^+^ (Table 3). By contrast, 300 *µ*M Zn^+^ rarely affected the inactivation of I8Q-H382Q (Fig. 9e, Table 1) as well as its recovery time course (Fig. 9f, Table 3). These results suggest that H382 is involved in C-type inactivation of AKv1, and that the Zn^2+^ sensitivity of C-type inactivation in AKv1 depends on the Zn^2+^-binding to H382 in the pore turret.

## Discussion

*Aplysia* Kv1 channel, AKv1, shows robust N-type inactivation which can be approximated by a single step process [14, 52, 53, 54]. Compared to the fast onset of inactivation the recovery from inactivation in low external K^+^ condition is extremely slow [14, 52], and the recovery rate from the inactivation is almost identical to that from C-type inactivation in the amino-terminal deletion mutant, suggesting that AKv1 usually recovers from C-type inactivated state [54] like mammalian Kv1.4 [56]. In either channel, the N-C coupling must be quite efficient compared to *Drosophila shaker* channel [28] in which a large component of the recovery from inactivation (*>*50–60%) seems to be directly from N-type inactivated state in low external K^+^ condition [4, 10].

As described in Introduction, N-type inactivation of Kv1 channel is actually multistep process [54, 66, 67, 73]. The amino-terminal domain of the channel (or Kv*β*) binds to the T1 domain at first before entering into the pore through the window beneath the transmembrane domain [20, 54, 73]. The extended amino-terminus then slips into the window and occludes the open pore [20, 66, 67, 73]. In general, the amino-terminal hydrophobic region is necessary for the final pore block and the following polar region is important for attaching to the T1 domain [27, 54, 66, 73]. N-type inactivation of AKv1 has been extensively studied using several mutants, and several subregions in the aminoterminal domain which are critical for multistep N-type inactivation have been identified [51, 53, 54]. In the N-type inactivation of AKv1, the amino-terminal five amino acids are important for the pore block and the following *∼*9 amino acids are important for the docking of amino-terminal region to the window above T1 domain [51]

Among the previously tested amino-terminal mutants in AKv1, we are interested in the mutant, I8Q [51]. In high K^+^ condition which inhibits C-type inactivation, the inactivation of I8Q is approximated by a single exponential with *τ_inacti_* comparable to that of AKv1, suggesting that the fast inactivation of I8Q is N-type [51]. In contrast to AKv1, however, a large stationary current is observed at the end of long depolarizing pulse in I8Q [51]. The fraction of the conducting channels at the end of long depolarizing pulse to +40 mV in high K^+^ condition is *∼*50% compared to *<*5% in AKv1, indicating that the inactivation of I8Q should be less stable. In a simple pore block model for N-type inactivation, the results can be easily explained if we assume the I8Q mutation increases the dissociation rate constant of the N-terminal blocking particle (*β*) with concomitant decrease of the association rate constant (*α*): In such case, *τ_inacti_* (i.e., 1/[*α*+*β*]) can be similar even if the stationary current which is proportional to *β*/[*α*+*β*] becomes larger. To characterize properties of I8Q, we compared the inactivation of I8Q in low and high K^+^ condition in the present study. The inactivation of I8Q in ND96 (2mM K^+^) was approximated by sum of two exponentials. The fast *τ_inacti_* of I8Q was similar to *τ_inacti_*of AKv1, suggesting again that the fast inactivation of I8Q is N-type. The appearance of slow component in the inactivation of I8Q in ND96 can reflect less stable pore block and less efficient N-C coupling (see below).

When the inactiavting prepulse was more than several hundread msec, the single exponential *τ_rec_* of AKv1 was the same to that of ΔN in ND96. In high K^+^, the *τ_rec_* of AKv1 (*∼*200 msec) was more than 10 times shorter than that in ND96 and the recovery from inactivation was *∼*9 times faster than that of ΔN (see Table 3 and 5). If we can assume C-type inactivation of AKv1 is well inhibited in high K^+^ condition [16, 35, 40, 54], these results are consistent with a notion that the recovery of AKv1 in high K^+^ is dominated by a pseudo single step recovery from N-type inactivated state [54]. When the inactivating prepulse was shorter than a few hundred msec, however, the recovery from inactivation of AKv1 in ND96 was better fitted with two exponentials: the fast *τ_rec_*was 500–700 msec and the slow *τ_rec_* was 3–4 sec (Table 2). Because 70–90% of the recovery was due to the slow process, the over-all recovery time course was still comparable to that of C-type inactivation in ΔN. These results are consistent with the current hypothesis that the N-C coupling in AKv1 is quite efficient and the recovery from inactivation is mostly from C-type inactivated state [54]. Because the fast *τ_rec_* observed after shorter inactivating prepulse in AKv1 was in a similar range to the *τ_rec_* in high K^+^, the fast component of recovery is likely to reflect the recovery from N-type inactivated state as assumed in *Drosophila Shaker* channel [4].

The recovery of I8Q in ND96 was always approximated by two exponentials and its time course was changed drastically depending on the prepulse length (Fig. 2d, Table 2). In high K^+^ condition, the recovery was dominated by the fast process (the slow component was *∼*13%, see Fig. 3b, Table 3). Again, the results can be explained by the inhibition of C-type inactivation by high K^+^, and the similarity between the recovery after the 50 msec prepulse in ND96 and the one after the 1 sec prepulse in high K^+^ (see Fig. 2d and Fig. 3b) suggests that the inactivated state of I8Q after the 50 msec prepulse in ND96 are mostly N-type inactivated state. The *τ_rec_* of AKv1 in high K^+^ was shorter than the fast *τ_rec_* of AKv1 in ND96 which was observed when the duration of inactivating prepulse was less than a few hundred msec. Also, the fast *τ_rec_* of I8Q in high K^+^ was shorter than that obtained in ND96 (see Table 3). These results can be explained at least partly by a push-off effect of external K^+^ on N-type inactivation at hyperpolarized recovery potential [10].

It might be interesting to compare the recovery data in the present experiments with the analysis of amino-terminal peptide (ShB-p) block in Sh-IR [46, 47]. In Sh-IR, ShB-p block of the channel can recover with two components at -120 mV: the fast *τ* is *<*1 msec and the slow one is *∼*90 msec in low K^+^ condition (2 mM) and *∼*23 msec in high K^+^ (140 mM) [46]. Although the slow *τ* in low K^+^ is a little bit slower than the fast *τ_rec_* of I8Q in ND96 (Table 3), the slow *τ* in high K^+^ is similar to the fast *τ_rec_* of I8Q in high K^+^. This similarity may suggest that the fast recovery from the inactivation in I8Q, especially in high K^+^ condition, is mechanistically the same to dislodging of the blocking peptide from the open pore. On the other hand, the *τ_rec_* of AKv1 in high K^+^ was still much longer than the slow *τ* of ShB-p block recovery in Sh-IR, implying that ”N-type inactivation” of AKv1 is somewhat more sticky.

Isoleucine at 8th from the amino-terminus of AKv1 is in a so called ”IP-region” which is supposed to bind the window above T1 domain before the final pore block in N-type mechanism [51]. The I8Q mutation probably deteriorates the binding affinity of ”IP- region” to the window above T1 domain and reduces a probability of the channel in the pre-block state, which would preclude a stable insertion of the amino-terminus into the pore. The importance of ”IP-region” is actually observed earlier by Hoshi et al (1991) in their first paper documenting N-type inactivation of *Drosophila Shaker* channel, in which the mutation of leucine (L7) in the amino-terminus of *Drosophila Shaker* channel is shown to inhibit N-type inactivation [27].

Although N-type inactivation and C-type inactivation are mechanistically distinct processes, the two mechanisms are coupled in the wild type Kv channels, which is first demonstrated in *Drosophila Shaker* channel [28]. Baukrowitz and Yellen (1995) have shown that external [K^+^] dependent cumulative inactivation of *Drosophila Shaker* channel is actually due to the accelerated C-type inactivation following N-type inactivation and the coupling between the two inactivation mechanisms has been elegantly explained by external K^+^ dependency of C-type inactivation [4]. In this hypothesis, a fast plugging of the open pore by the amino-terminal structure efficiently stops the outflow of K^+^ and reduces the local [K^+^] near the external mouth of the pore, which accelerates C-type inactivation [4]. Another likely hypothesis for the N-C coupling is that the binding of amino-terminal structure to the pore allosterically changes the structure involved in C-type mechanism, which speeds up C-type inactivation [36, 44, 56]. The two hypothetical mechanisms are not mutually exclusive and may well coexist: ex., the hypothetical allosteric conformational change leading to C-type inactivation may be sensitive to external K^+^.

One of the prominent feature of AKv1 is the cumulative inactivation by short repetitive pulses [14], which has strong influence on the adaptation of spiking activity of *Aplysia* neurons [41]. Functionally similar cumulative inactivation has been analyzed in other Kv channels with or without N-type mechanism [1, 2, 4, 5, 42]. The cumulative inactivation of *Drosophila Shaker* channel which is a prototype Kv channel showing N-type inactivation has been explained by the external [K^+^] dependent acceleration of C-type mechanism in the N-C coupling [4] (see above). Also, C-type inactivation is considered to be essential for the cumulative inactivation in Kv1.4 which also shows a robust N-type inactivation [5]. Although the cumulative inactivation of AKv1 is partly K^+^ sensitive and is likely due to the combination of N-type and C-type mechanisms [16, 60], we think a key determinant of the cumulative inactivation in AKv1 is a stability of the pre-block state in N-type inactivation as discussed below.

In the present study, we examined the cumulative inactivation by tandem short pulses (+40 mV, 20 msec) using a wide range of inter-pulse potentials in ND96 and high K^+^ solution. We found that the inactivation can proceed even at –80 mV in ND96, and that the activatable channels 5 msec after the 1st pulse were reduced to *∼*60% irrespective of the inter-pulse potential or external [K^+^]. Similar tendency was also observed in I8Q but the cumulative inactivation was much less. Because the inactivation of AKv1 as well as I8Q during a 20 msec depolarizing pulse is limited and similar, the change in local [K^+^] near the external mouth of the pore by K^+^-efflux is not considered to be different, discounting the hypothesis that the difference in local [K^+^] is a main cause of different cumulative inactivation between AKv1 and I8Q.

We also observed clear gating charge immobilization in AKv1-W391F but not in I8Q- W391F or ΔN-W391F by using the two pulse protocol, which is correlated to the cumulative inactivation of AKv1. Because a 10 msec depolarizing pulse used in the gating current experiments is too short to see meaningful N-type inactivation in the ionic currents, the charge immobilization is not likely due to the pore block by N-type mechanism during the 1st pulse. Rather, we think the charge immobilization observed in AKv1-W391F is related to the occupancy of the pre-block state during the 1st pulse. During the inter-pulse interval at hyperpolarized potential, some of the channels in the pre-block state deactivate but others must quickly enter the pore blocked state, which inhibits the recovery of gating charge and increases a chance to enter C-type inactivated state. Alternatively, the pre-block state itself may preclude the recovery of gating charge. The gating charge immobilization is much less obvious in I8Q because I8Q is less well to occupy the pre-block state as discussed above. We think the above mentioned evidences favor an idea that the stability of the pre-block state in N-type inactivation but not external [K^+^] is essential for the cumulative inactivation of AKv1.

External Zn^2+^ inhibited the recovery from inactivation of AKv1, suggesting that Zn^2+^ potentiates C-type inactivation of AKv1. A specific action of Zn^2+^ on C-type inactivation is more clearly observed in I8Q. The slow component of inactivation in I8Q was specifically accelerated by Zn^2+^, and the recovery time course of I8Q in 300 *µ*M Zn^2+^ became a slow single exponential, *τ_rec_*of which was close to that of AKv1. Although its potency was slightly reduced, Zn^2+^ accelerated C-type inactivation of ΔN even in high K^+^ condition, suggesting that the effects of K^+^ and Zn^2+^ on C-type inactivation are not mutually exclusive. Although the inactivation of *Drosophila Shaker* channel is not affected by Zn^2+^ [61, 70], Zn^2+^ is known to potentiate the inactivation of some Kv1 channels. The recovery from inactivation in Kv1.4-Kv1.1 heteromer which is supposed to be endogenous Kv channels in hippocampal neurons is hindered by Zn^2+^ [29]. Zn^2+^ apparently blocks Kv1.5 and the effect appears to be due to the stabilization of C-type inactivation [31, 72]. The Zn^2+^-induced block of Kv1.5 is, however, diminished by external K^+^ as low as a few mM [72]. The sensitivity of Zn^2+^-block in Kv1.5 to external K^+^ is contrast to the result in ΔN, and implies a difference in C-type mechanism between the two channels.

A378E and D379P show N-type inactivation indistinguishable with AKv1 but impaired C-type mechanism [60]. In the present study, these two mutants showed much faster recovery from inactivation than AKv1, *τ_rec_* of which was not affected by Zn^2+^. The results are consistent with the hypothesis that Zn^2+^ specifically acts on the C-type mechanism of AKv1. In consistent with this, the effect of Zn^2+^ was much diminished in ΔN-A378E. However, the inactivation of ΔN-A378E was rather enhanced by high K^+^ and the overall time course of the recovery from inactivation was not affected by high K^+^. Because the enhanced inactivation by high K^+^ is a characteristic of U-type inactivation in some Kv channels [33, 34], the A378 mutation appears to change the property of C-type inactivation substantially. Indeed, the effects of K^+^ and Zn^2+^ on *τ_rec_* of I8Q-A378E were still observed although the inactivation of I8Q-A378E became less complete compared to I8Q. The A378E mutation (and probably the D379P mutation), therefore, does not seem to inhibit Zn^2+^-binding specifically.

A more clear picture was obtained by another pore turret mutant, H382Q. C-type inactivation of ΔN-H382Q was slower than that of ΔN and was sped up in high K^+^ solution, indicating some functional similarity of this mutant to ΔN-A378E. However, Zn^2+^ had essentially no effect on the slow inactivation of ΔN-H382Q and the recovery from inactivation was little affected by either Zn^2+^ or high K^+^. On the other hand, the inactivation of I8Q-H382Q was essentially similar to that of I8Q and the recovery from inactivation was K^+^-sensitive. External Zn^2+^ did not affect the inactivation of I8Q-H382Q as well as the recovery time course at all, suggesting that H382 is a key residue for the effect of Zn^2+^ on C-type inactivation in AKv1. Taken together, these results are consistent with a notion that external Zn^2+^ specifically accelerates C-type inactivation of AKv1 by binding to a site involving H382 in the pore turret. The different K^+^-sensitivity between ΔN-H382Q and I8Q-H382Q (or between ΔN-A378E and I8Q-A378E) indicates that the C-type mechanism of the amino-terminal deletion mutant is not entirely the same to that in the N-C coupled inactivation, which favors the allosteric conformational change hypothesis for the N-C coupling [36, 44, 56].

In summary, we compared the inactivation between AKv1 and I8Q under several conditions. We found that the N-C coupling which is quite efficient in AKv1 is functional but inefficient in I8Q. By utilizing I8Q and its mutants, we showed rather clearly that Zn^2+^ as well as high K^+^ specifically affect C-type inactivation, and that H382 in the pore turret is involved in Zn^2+^ binding. Because of the temporally dissociated but still coupled N-type and C-type inactivation, the I8Q mutant should be a useful model for the exploration of molecular mechanism of the N-C coupling, which is an important determinant of the frequency dependent spike accommodation of neurons.

## Declarations

### Authors’ contributions

YF conceived research; YF designed experiments; TI, KI and YF performed experiments; TI, KI and YF analyzed data; YF wrote a manuscript; TI, KI and YF approved the final version of the manuscript.

### Availability of data and materials

The data support the findings of this study are available from the corresponding author upon reasonable request.

