## Supplemental Tables for "Stability of N-type inactivation and the coupling between N-type and C-type inactivation in the *Aplysia* Kv1 channel"

Supplementary tables

Table 1
Table 2

Table 1. Cumulative inactivation assessed by exponential fittings

| | | AKv1 | | | | I8Q | | | |
|---|---|---|---|---|---|---|---|---|---|
| solution | ip | A1 | $\tau1$ | A2 | $\tau2$ | A1 | $\tau1$ | A2 | $\tau2$ |
| ND96 | -40 | 0.36±0.10 | 8.9±1.7 | -0.69±0.03 | 4201.6±639.8 | 0.19±0.05 | 6.1±0.9 | -0.30±0.06 | 209.6±29.3 |
| ND96 | -80 | 0.14±0.03 | 3.9±1.2 | -0.53±0.06 | 1476.9±87.3 | - | - | -0.22±0.05 | 78.83±4.5 |
| ND96 | -120 | - | - | -0.45±0.07 | 602.2±53.1 | - | - | -0.19±0.04 | 40.9±0.8 |
| ND96 | -160 | - | - | -0.41±0.08 | 262.6±33.1 | - | - | -0.15±0.04 | 31.6±1.9 |
| high $K^+$ | -40 | 0.25±0.06 | 5.6±2.0 | -0.57±0.04 | 1037.7±184.8 | 0.19±0.02 | 5.7±0.8 | -0.26±0.04 | 72.4±13.1 |
| high $K^+$ | -80 | - | - | -0.46±0.07 | 271.1±26.3 | - | - | -0.19±0.03 | 29.7±2.2 |
| high $K^+$ | -120 | - | - | -0.42±0.08 | 80.8±5.9 | - | - | -0.18±0.03 | 12.8±1.3 |
| high $K^+$ | -160 | - | - | -0.39±0.08 | 29.4±2.0 | - | - | -0.17±0.03 | 5.4±0.4 |

Mean±SD of parameters estimated by exponential fitting of the relationship between $I_{2nd}/I_{1st}$ and the inter-pulse interval are shown (n=5).

ip: inter-pulse potential (mV)

A1, A2: the relative amplitudes of exponential components

$\tau1$, $\tau2$: time constants (msec) of exponential components

Relative amplitude of maximum recovery (Y0) was assigned to 1

Table 2. Exponential recovery of the ON gating charge

| channel | ip | A1 | $\tau$1 | Y0 | n |
|---|---|---|---|---|---|
| AKv1 | -80 | -0.463±0.057 | 31.4±5.4 | 0.572±0.060 | 9 |
| | -100 | -0.635±0.067 | 17.2±3.0 | 0.744±0.048 | 9 |
| | -120 | -0.754±0.084 | 8.5±1.8 | 0.838±0.055 | 9 |
| | -140 | -0.846±0.092 | 4.1±1.0 | 0.868±0.059 | 9 |
| I8Q | -80 | -0.820±0.118 | 31.2±2.6 | 0.905±0.060 | 4 |
| | -100 | -0.915±0.081 | 15.8±1.6 | 0.972±0.026 | 4 |
| | -120 | -0.982±0.092 | 7.9±1.5 | 1.011±0.041 | 4 |
| | -140 | -1.081±0.123 | 4.0±0.8 | 1.024±0.057 | 4 |
| ΔN | -80 | -0.973±0.029 | 32.1±1.1 | 1.005±0.028 | 3 |
| | -100 | -0.999±0.031 | 15.3±0.8 | 1.014±0.031 | 3 |
| | -120 | -0.987±0.035 | 6.8±0.5 | 0.994±0.034 | 3 |
| | -140 | -0.998±0.032 | 3.5±0.6 | 1.002±0.032 | 3 |

Mean±SD of parameters estimated by a single exponential fitting (see Materials and methods) of the relationship between the relative amplitude of the gating charge at 2nd pulse (Q2/Q1) and the inter-pulse interval are shown.

ip: inter-pulse potential (mV)

A1: relative amplitude of an exponential component

$\tau$1: time constant (msec)

Y0: relative amplitude of an estimated maximum recovery

n: the number of tested oocytes
